# Single cell 3’UTR analysis identifies changes in alternative polyadenylation throughout neuronal differentiation and in autism

**DOI:** 10.1101/2020.08.12.247627

**Authors:** Manuel Göpferich, Nikhil Oommen George, Ana Domingo Muelas, Alex Bizyn, Rosa Pascual, Daria Fijalkowska, Georgios Kalamakis, Ulrike Müller, Jeroen Krijgsveld, Raul Mendez, Isabel Fariñas, Wolfgang Huber, Simon Anders, Ana Martin-Villalba

## Abstract

Autism spectrum disorder (ASD) is a neurodevelopmental disease affecting social behavior. Many of the high-confident ASD risk genes relate to mRNA translation. Specifically, many of these genes are involved in regulation of gene expression for subcellular compartmentalization of proteins^1^. Cis-regulatory motifs that often localize to 3’- and 5’-untranslated regions (UTRs) offer an additional path for posttranscriptional control of gene expression. Alternative cleavage and polyadenylation (APA) affect 3’UTR length thereby influencing the presence or absence of regulatory elements. However, APA has not yet been addressed in the context of neurodevelopmental disorders. Here we used single cell 3’end sequencing to examine changes in 3’UTRs along the differentiation from neural stem cells (NSCs) to neuroblasts within the adult brain. We identified many APA events in genes involved in neurodevelopment, many of them being high confidence ASD risk genes. Further, analysis of 3’UTR lengths in single cells from ASD and healthy individuals detected longer 3’UTRs in ASD patients. Motif analysis of modulated 3’UTRs in the mouse adult neurogenic lineage and ASD-patients revealed enrichment of the cytoplasmic and polyadenylation element (CPE). This motif is bound by CPE binding protein 4 (CPEB4). In human and mouse data sets we observed co-regulation of CPEB4 and the CPEB-binding synaptic adhesion molecule amyloid beta precursor-like protein 1 (APLP1). We show that mice deficient in APLP1 show aberrant regulation of APA, decreased number of neural stem cells, and autistic-like traits. Our findings indicate that APA is used for control of gene expression along neuronal differentiation and is altered in ASD patients.

## INTRODUCTION

The brain entails over a million of highly specialized neurons that are generated throughout embryonic development from neuroepithelial and radial glia cells. A set of these radial glia cells (neural stem cells: NSCs) is retained in the adult brain. The largest known repertoire of NSCs locates to the walls of the lateral ventricles in the ventricular-subventricular zone (vSVZ). NSCs within the vSVZ transition through different states of activation to eventually produce a neurogenic progeny. Whereas in humans this progeny is found in the striatum^2^ in the mouse, it matures into olfactory bulb interneurons and integrates into the olfactory bulb network. The process of generation of olfactory bulb neurons in rodents offers a simple system to study neuronal maturation and therefore potentially provides insights into neurodevelopmental disorders.

Generation of highly specialized neurons involves posttranscriptional regulation of gene expression^3,4^. An important aspect of such regulation is alternative cleavage and polyadenylation (APA), which affects around two-thirds of all mammalian genes^5–8^. Through APA one among several polyadenylation signals (PAS) of an mRNA is selected, thereby influencing the length of the 3’UTR. This alternative usage of PAS allows for inclusion or exclusion of various regulatory elements, which can influence the stability, translation efficiency and cellular localization of transcripts^6–8^. APA has been implicated in various cellular processes such as cell proliferation^9,10^, cell fate regulation^11^, and senescence^1^, but also diseases like cancer^6^. 3’UTRs tend to be longer in neurons than in other cell types of the brain^8,12,13^ and its involvement in neurodevelopmental disorders have been shown for individual genes and hypothesized as potential player in Autism Spectrum Disorder (ASD)^14,15^.

## RESULTS & DISCUSSION

To study a potential role of APA for stem cell activation and further differentiation into a neuron we analyzed the single cell transcriptome of the olfactory bulb lineage comprising quiescent and active NSCs, transient amplifying progenitors and neuroblasts isolated from the vSVZ of young and old mice (2 and 22 month old)^16^. While the purpose of our previous study was to compare these two pools to find age-dependent effects, we now ignore the age information and treat them as replicates, i.e., all results reported here are valid independent of age. In this data set, each cell has been assigned a “pseudotime” value that marks its position along the trajectory from quiescent NSCs to neuroblasts (Fig. 1a). The cells’ transcriptome was sequenced by 10X Chromium single-cell RNA-Seq, which uses poly-dT primers to collect mRNA and thus allows pinpointing the end for each transcript molecule. These 3’ ends fall onto sharply defined peaks (Fig. 1b, left panel, Extended Data Fig. 1a-b), which are typically ∼20 bp downstream of the canonical polyadenylation signal, the hexamer AAUAAA (Fig. 1c), and also agree well with the Ensembl annotation for 3’ gene ends (Extended Data Fig. 1c). For each transcript molecule we calculated the length of its 3’UTR and examined for each gene if the selection of alternative PAS changes throughout the neurogenic lineage at single cell resolution. For many genes, we observed such changes. For example, the gene Pea15a shifts its usage to the proximal PAS (Fig. 1b). We quantify these changes by calculating for each gene the correlation coefficient between 3’ UTR length (average for each cell) and the cell’s pseudotime (Fig. 1d). A positive or negative coefficient indicates lengthening or shortening of the 3’UTR with maturation, respectively. We found good agreement between the two samples despite their age, whenever the correlation was nominally significant (Fig. 1e). However, correlations cannot capture complex changes like, for example, for Sox4 (Extended Data Fig. 1d), where the usage of the middle PAS increases with lineage progression while that of proximal and distal PASs decreases. Hence, we examined the fractional usage of each PAS over pseudotime by assigning reads to PAS, as most reads fell into well-defined poly(A) sites. Thereafter, we performed multinomial spline regression to get smooth usage fraction curves (Fig. 1f lower panel, Extended Data Fig. 1a-b, right panel). This regression yields a score that quantifies how strongly poly(A)-site-selection depends on pseudotime. These regression scores agreed well between both biological samples (Extended Data Fig. 1e). Our analysis unveiled a dynamic regulation of various genes by poly(A)-site selection along the NSC lineage progression. Genes with the GO term ‘central nervous system development’ exhibit high regression scores for APA (GO term analysis in Extended Data Fig. 1f). Notably, these genes were also enriched in categories associated with neurodevelopmental disorders using two independent disease-related databases (Fig. 1g). Specifically, autistic spectrum disorder was enriched in both databases. Altogether, this analysis indicates that neuronal differentiation involves APA changes in neurodevelopmental genes that are dysregulated in ASD.

**Figure 1.**
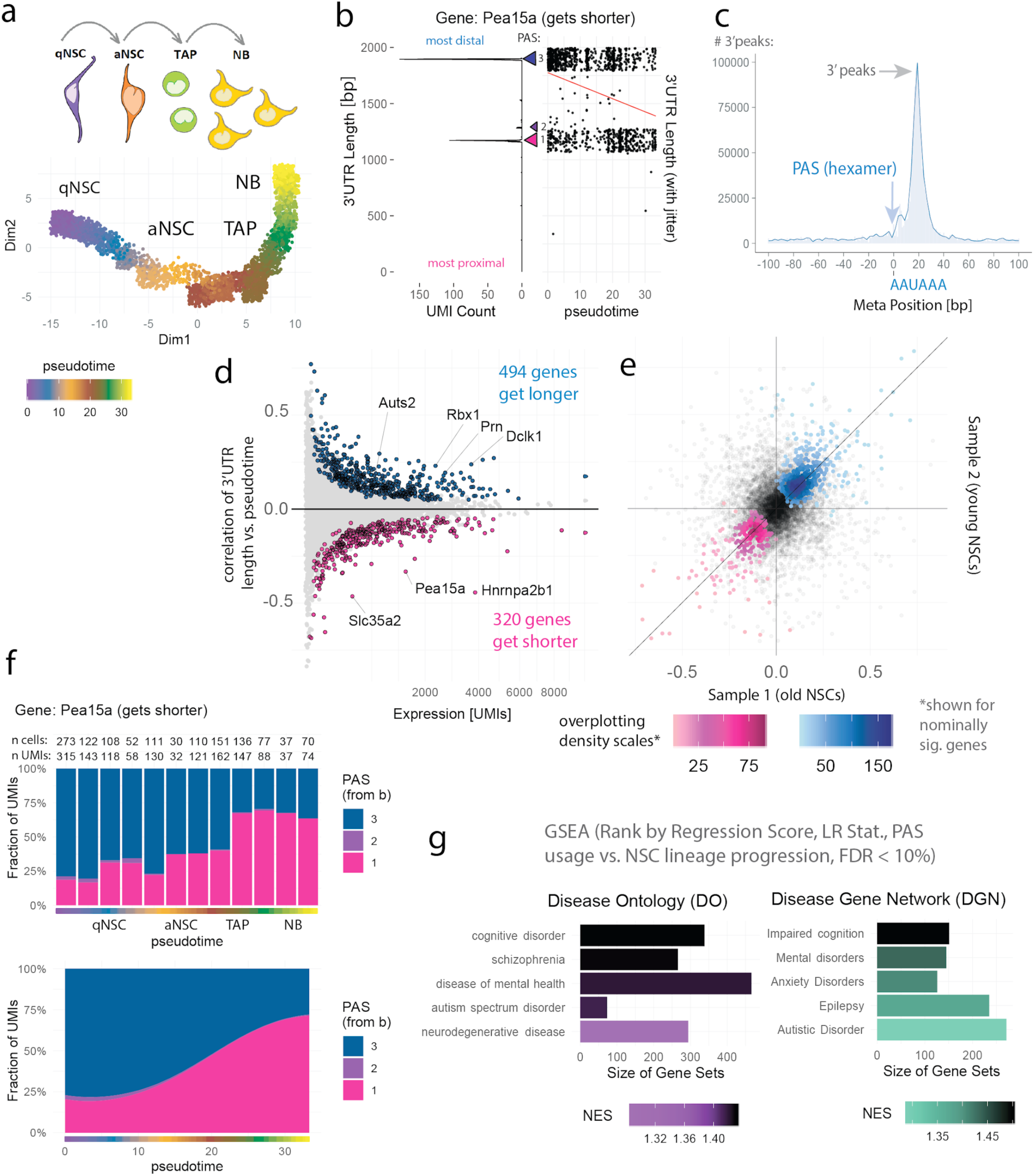
Alternative polyadenylation changes with NSC lineage progression and affects neurodevelopmental risk genes. **a**, scRNA-seq of NSCs (*Kalamakis et al. 2019*) differentiation to NBs, ordering in pseudotime (purple/blue: qNSCs, orange/brown: aNSCs, green: TAPs, yellow: neuroblasts) representing lineage progression, upper subpanel: schematic representation of the lineage, lower sub-panel: two-dimensional representation created with *monocle2*. **b**, Example gene Pea15a (phosphoprotein enriched in astrocytes 15A, also known as Mat1) getting shorter with lineage progression, left sub-panel: UMI counts and their respective 3’ mapping positions. (summed over cells), right sub-panel: mean mapping pos. per cell (y axis) versus pseudotime as in a (x axis), linear regression line (in red). **c**, Density profile of 3’ mapping positions, relative to the AAUAAA hexamer, shown as meta-gene analysis *i*.*e*. summed for expressed genes. **d**, For every gene with multiple PAS (APA genes): summed expression (x-axis) against correlation of mean 3’UTR length per cell vs. pseudotime (y-axis), 3’UTR shortening or lengthening marked as magenta and blue points respectively. **e**, Correlations as in d, computed independently for both biological replicates as scatter plot, colored points indicate nominally sig. genes, separate color scales for shortening (magenta) and lengthening (blue). **f**, Upper panel: fraction of different PASs from b per pseudotime window for Pea15a (blue = distal, long 3’UTR, magenta = proximal, short 3’UTR), lower panel: approximation of changes in 3’UTR usage over pseudotime utilizing multinomial regression splines. **g**, Gene set enrichment analysis (GSEA), genes ranked by the strength of 3’UTR changes with lineage, genes from d translated into human gene orthologs, a high normalized enrichment score (NES) indicates association of disease and 3’UTR changes.

To address whether there are changes in 3’UTR length in humans diagnosed with ASD we used NucSeq RNA profiles of single neurons from a cohort of autistic and control individuals^17^ (Fig. 2a, quality control in Extended Data Fig. 2a). To dissect this effect, we computed changes in 3’UTR length comparing ASD to controls for each cell type, respectively (Fig. 2a, Methods, ANOVA). Across all cell types we observed a global lengthening of 3’UTRs in autism patients in comparison to controls (Fig. 2b). This is shown in detail for layer 2/3 excitatory neurons (Fig. 2c) and for SOX4 and NCAM2 that are longer in autism compared to controls (Fig. 2d). This effect can also be visualized for other cell types; VIP interneurons (IN-VIP) and fibrous astrocytes (AST-FB) (Extended Data Fig. 2b-c). Gene set enrichment analysis of genes ranked by the 3’UTR length changes (autism vs. control) revealed that neurotrophin signaling (NTRKs), axon outgrowth, cell cycle and mRNA translation were among the categories with the highest enrichments (Fig. 2e). Defective axonal pathfinding and synaptic function lead to aberrant brain connectivity and dysfunction that account for repetitive behaviors and impaired social interaction: the core symptoms of ASD. Accordingly, ASD risk genes encode proteins either directly involved in synaptic function and connectivity such as neurotransmitters or ion channels, or involved in regulation of mRNA transport and translation^18^. The list of identified translation regulators encompasses components of the translation machinery, miRNAs and RNA binding proteins. Both miRNAs and RNA binding proteins exert their function through binding to motifs within the 5’ and 3’-UTR of the mRNAs. Despite the obvious involvement of the 3’UTR in RNA processing, this is the first study discovering global changes in 3’UTR length in ASD patients based on 3’ scRNAseq^c.f.14^.

**Figure 2.**
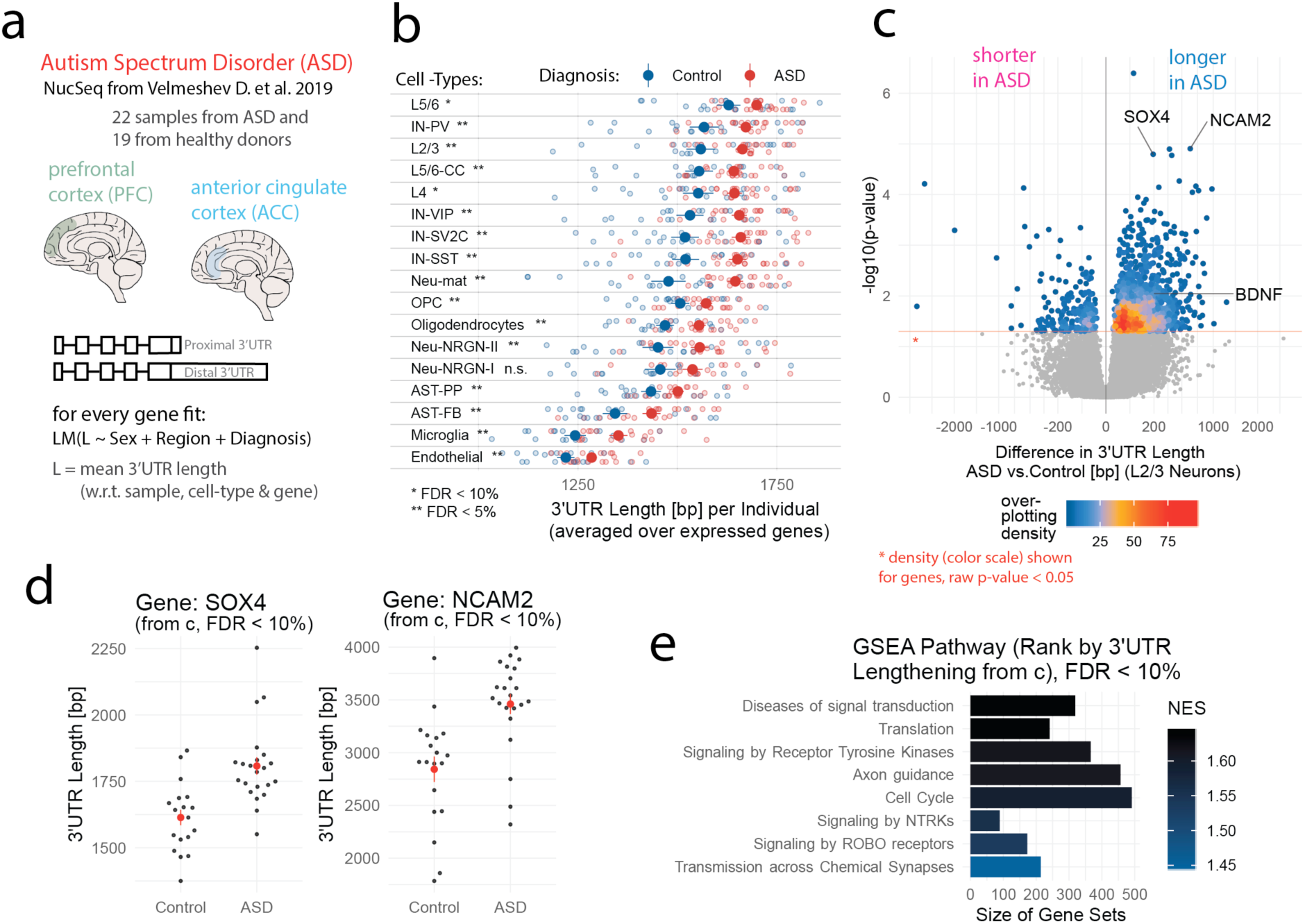
ASD patients show 3’UTR lengthening compared to healthy individuals. **a**, Schematic depiction of analysis strategy, test for 3’UTR alterations in an autism cohort vs. healthy individuals from Velmeshev et al. 2019, test with linear models (LM) 3’UTR mapping positions considering per gene and cell-type: brain region, sex and diagnosis. **b**, Meta-gene analysis of 3’UTR length shows a lengthening trend in autism, every point, mean 3’UTR length averaged over all expressed genes for one sample and one cell type (red for ASD, blue for healthy control, big points show group means), multiple t-tests for each cell-type with FDR control, two-sided (Benjamini-Hochberg). **c**, Volcano plot showing 3’UTR alterations in layer 2/3 excitatory neurons comparing ASD patients to controls for each gene, 3’UTR length differences (x-axis), p-value estimated from linear models (from a, y-axis), over-plotting color scale applied to genes with uncorrected p-values < 0.05 to show the overall trend. **d**, Example genes from c, every point average 3’UTR length over layer 2/3 excitatory neurons per sample, red points group means, linear model (ANOVA) from a, left, SOX4 (SRY-box transcription factor 4), right, NCAM2 (neural cell adhesion molecule 2), both genes get longer in ASD vs. healthy control. **e**, Gene set enrichment analysis, genes ranked by the genes getting longer im ASD vs control patients from c (fold-changes), a high normalized enrichment score (NES) indicates association of 3’UTR lengthening in ASD patients with the respective pathway.

Examination of neuronal differentiation in the mouse olfactory bulb lineage and in neurons from ASD patients revealed changes in APA, however the impact of these changes is unclear. To see how activation-driven changes in APA selection impact translation, we turned to an in-vitro system in order to have enough material for proteomics profiles. We treated NSC cultures with BMP4 or EGF in order to model, respectively, quiescent and active NSCs (Fig. 3a). Transition to quiescence was verified by a strong reduction of the proliferation marker Mki67 in BMP4-treated NSCs as compared to their EGF-treated counterparts (Extended Data Fig. 3a). Single cell RNAseq of this *in vitro* system revealed that many genes exhibited comparable expression levels between *in vitro* cultures and their *in vivo counterparts* (directly isolated from the brain) (Fig. 3b and Extended Data Fig. 3b-c). We next assessed to which extent 3’UTR changes are conserved between cultures and *in vivo* NSCs. For this we computed for every PAS the log2-fold-change of its usage between aNSCs and qNSCs *in vitro* and *in vivo*. Both data sets showed substantial correlation (R = 0.5). We therefore conclude that a substantial amount of 3’UTR changes can be reproduced *in vitro*. Interestingly, gene categories that deviate in their 3’UTR usage between *in vivo* and *in vitro* relate to responses to the environment hinting at the absence of environmental signals in the cultures that would be needed to trigger APA of these specific genes (Extended Data Fig. 3d). Subsequently, we isolated proteins from NSCs cultures, and identified and quantified them by mass spectrometry (Fig. 3c, quality control in Extended Data Fig. 3e). Many of the changes in mRNA levels from qNCSs to aNSCs are mirrored in changes in protein abundance (R = 0.7, Extended Data Fig. 3f). To evaluate differences between aNSCs and qNSC in mRNA translation we defined a translation index (TI), per gene: dividing protein abundance by mRNA levels. For genes undergoing 3’UTR shortening from qNSCs to aNSCs we observed a trend for higher protein outcome in aNSCs (Fig. 3d), which was consistent across biological replicates (Extended Data Fig. 3g). Hence, we conclude that 3’UTR shortening can increase protein production in the first transition from qNSC to aNSC, something that can be explained by missing repressive elements in the shorter 3’UTRs^6,9,19^.

**Figure 3.**
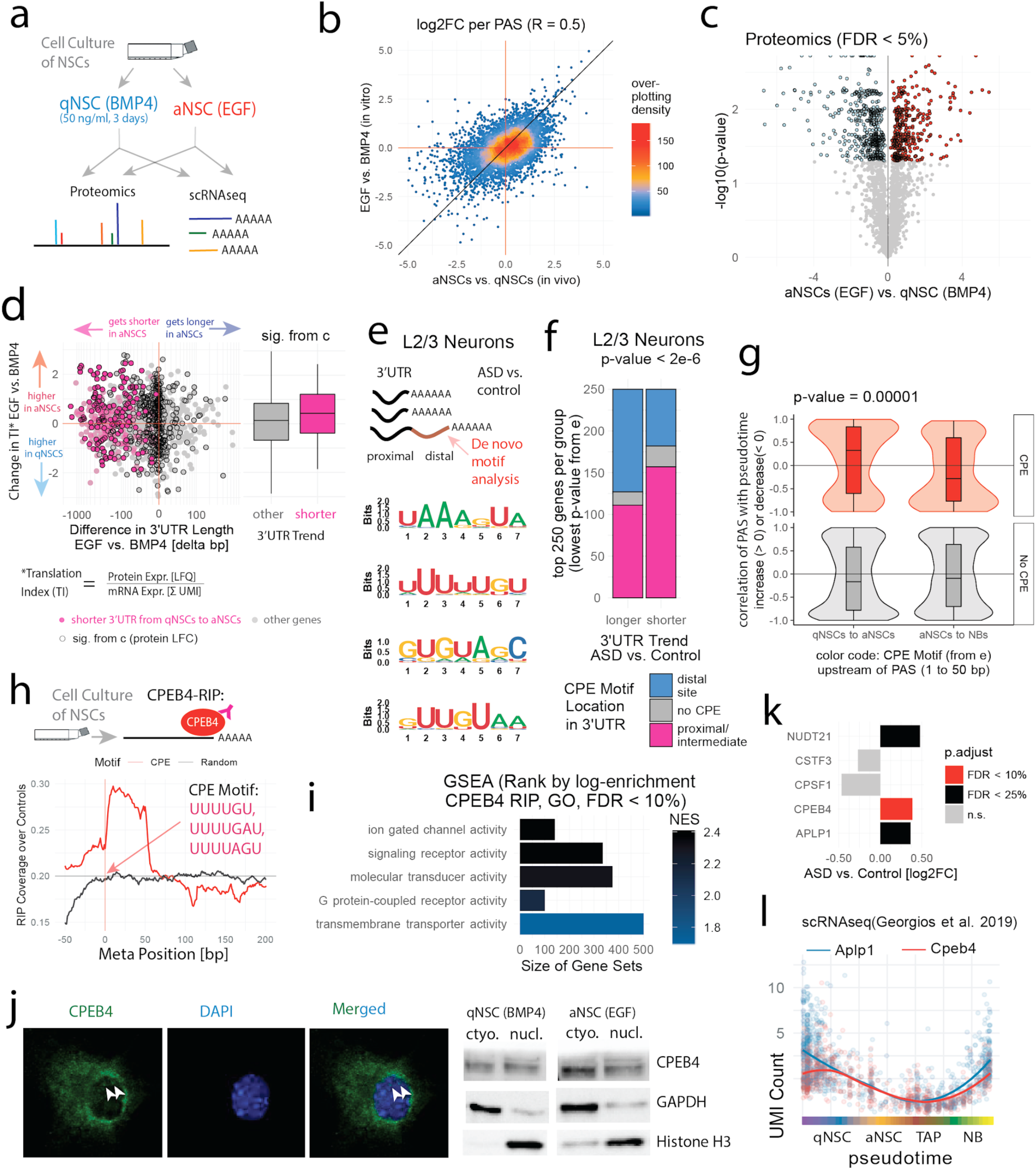
3’UTR shortening in NSCs results in increased protein outcome and binding of CPEB4 is predictive for the directionality of 3’UTR alterations with lineage progression. **a**, Schematic representation of experimental layout: culturing of NSCs, treated with BMP4 or EGF, for each treatment: scRNA-seq (mRNA and PAS detection) and total protein quantification by mass spectrometry, **b**, Log fold-change, every point represents one PAS (summed over cells, positive LFCs= higher PAS usage in aNSCs, negative LFCs = higher PAS usage in qNSCs), x-axis as reference (*in vivo* NSCs from Fig. 1), y-axis for *in vitro* NSCs. **c**, Volcano plot, differential protein abundance *in vitro* between aNSCs (EGF) and qNSCs (BMP4), proteins marked in red and blue (FDR < 5%, Perseus tool). **d**, Scatterplot, every point is a gene, difference in 3’UTR length aNSCs (EGF) and qNSCs (BMP4) on the x-axis vs. difference in translation index (TI: LFQ values by UMI counts) on the y-axis, boxplot on the left: comparison of other genes to 3’UTR shortening genes (c.f. Extended Fig. 3f-g). **e**, De-novo motif analysis utilizing the *homer2* tool (Methods), motifs enriched in distal 3’UTRs getting longer in ASD vs. controls compared to 3’UTRs getting shorter in ASD vs. controls. **f**, Association of 3’UTR alterations is ASD to CPEB4 binding, per gene classification whether the CPE motif from **h**, is in the distal site (human genes), proximal/intermediate or not present, shown per top 250 lengthening and shortening genes, respectively from Figure 2c, Chi-square test. **g**, Comparison of PAS containing the CPE motif (red color, motif from **f**) to other PAS (statistical control) depending on how the individual PAS usage changes from qNSC to aNSCs (left group) and aNSCs to NBs (right group), trend estimated by spline regression, ANOVA interaction test, two-sided. **h**, Signal from RNA immunoprecipitation (RIP) in cultured NSCs with CPEB4 antibody for CPE motif, mean coverage over RIP controls as meta-position, random position as statistical control shown as grey line. **i**, Gene set enrichment analysis, ranked by strength of CPEB4 binding to mRNAs, estimated as fold-change (RIP assay from h), a high normalized enrichment score (NES) indicates an overrepresentation of CPEB4 binding for respective gene categories. **j**, Detection of CPEB4 protein in the nucleus as shown by in situ stainings of *in vitro* NSCs; (left panel) and western blot of nuclear extracts (right panel; low GAPDH and high Histone H3, see also Supplementary Image. 1). The representative slice of a confocal stack shows CPEB4 speckles (white arrows) in the nucleus (DAPI stained). **k**, Differential expression of candidate genes in ASD vs. healthy controls, fold-changes from DESeq, x-axis, log2-fold-change in gene expression in L2/3 neurons, color code, p-value thresholds (FDR corrected). **l**, Co-expression of Aplp1 and Cpeb4 along the NSC lineage, each point indicates a single cell regression line fitted for both genes to show the trend.

As regulation of 3’UTR length controls the presence of motifs involved in posttranscriptional regulation of gene expression, we looked for a potential enrichment of regulatory elements in the changing 3’UTRs using homer2^20^ first in the human data set and then in the mouse data set. First, motif analysis comparing the distal region of the 3’UTR in top lengthening vs. top shortening genes in layer 2/3 excitatory neurons revealed an enrichment for the cytoplasmic polyadenylation element^21–23^ (CPE) in genes getting longer in ASD (Fig. 3e-f). CPE motifs have been identified in several dendritically localized neuronal transcripts^24,25^. Their binding to CPE binding proteins^26,27^ (CPEBs) regulates transport and translation of these transcripts. Specifically, CPEB1 and CPEB4 regulate poly-A tail length and thereby translation^28^. Furthermore, CPEB1 also acts in the nucleus, there affecting APA^28^. Of note, CPEB4 was recently identified as regulator of expression of high-confidence ASD risk genes in human and mouse neurons^29^. However, that study examined regulation of translation of ASD risk genes through poly(A)-tail lengthening but did not address a potential involvement of APA. We thus used our mouse data set to address this issue. De-novo motif analysis using homer2 of 3’UTR regions changing along neuronal maturation also unveiled an enrichment of CPE-like motifs (Extended Data Fig. 3d). Subsequently, we asked whether the CPE motif also influences PAS selection along neuronal differentiation. Selection of PAS flanked by an upstream CPE motif tended to increase upon activation (qNSCs to aNSCs) and to decrease upon differentiation (aNSCs to NBs) (Fig. 3g). We then examined if the 3’UTRs containing CPE-like motifs would indeed bind CPEB4. To this end, CPEB4-binding transcripts were immunoprecipitated (RIP) with an antibody against CPEB4 in cultured aNSCs and subsequently sequenced (Extended Data Fig. 4a-b). We detected over 900 mRNAs bound by CPEB4 (Extended Data Fig. 4b). These transcripts show enrichment for receptors, protein transporters and ion channel categories (Fig. 3i). The sequenced RNA binding site from the CPEB4-RIP assay highly correlated with the *in silico* predicted CPE motifs (meta-gene-analysis in Fig. 3h). Thereafter examination of expression levels of CPEBs revealed that CPEB4 is higher expressed along the neuronal lineage than the other CPEBs, with highest levels in qNSCs and neuroblasts (Extended Data Fig. 4c). Further, immunofluorescence staining as well as Western blotting revealed that CPEB4 is localized to the nucleus and the cytoplasm (Fig. 3j). Also, we found that the genes that get longer with lineage progression tend to have the CPE motif located rather in the distal PAS compared to the ones that get shorter (Extended Data Fig. 4e). Accordingly, protein expression of the transcripts binding to CPEB4 was higher in qNSCs than in aNSCs (Extended Data Fig. 4f). In summary, in human neurons and the rodent neurogenic lineage, changing 3’UTRs exhibit an enrichment of CPE. We could show that this motif is bound by CPEB4 in rodents, which suggest that not only CPEB1 but also CPEB4 can influence PAS selection.

**Figure 4.**
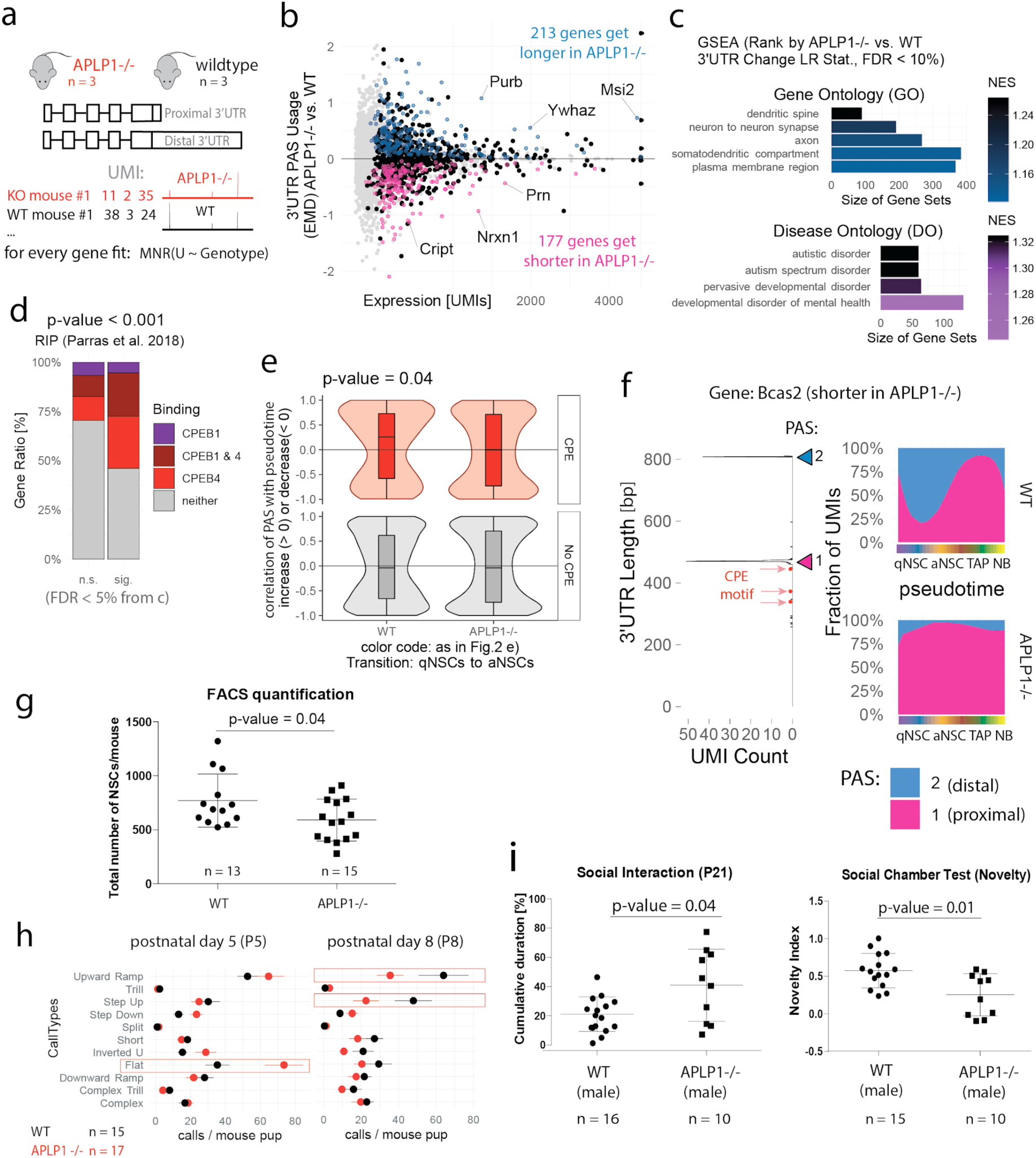
3’UTR alterations in APLP1-/- NSCs compared to wildtype bear a signature of CPEB4 binding and APLP1-/- mice show defects in the maintenance of the V-SVZ-NSC pool and autism-like traits. **a**, Schematic depiction of experimental layout, test for differential 3’UTR usage APLP1-/- vs. wildtype mice using scRNA-seq, single cells assigned to biological replicates by hash tag oligos (Ext. Fig. 4b). **b**, Differential 3’UTR usage (in NSCs, TAPs and NBs) comparing APLP1-/- to wildtype control, total expression summed over single cells (x-axis) vs. earth mover’s distance (EMD, y-axis) to assess directionality of 3’UTR changes, non-grey points = FDR < 5% (multinomial regression), magenta or blue if shorter or longer. **c**, Gene set enrichment analysis, ranked by strength of 3’UTR changes between APLP1-/- and wildtype (by multinomial regression from c), normalized enrichment score (NES), upper sub-panel, gene ontology, lower panel disease ontology (for respective human gene orthologs). **d**, Intersection of genes sig. from **c** with CPEB1 and CPEB4 substrates by Parras *et al*. 2018, color code: gene bound by CPEB1, CPEB4, by both or neither of both, genes marked as significant (sig.) or non-significant (n.s.) from **b** (APLP1-/- vs. wildtype 3’UTR changes), Chi square test. **e**, Comparison of PAS containing the CPE motif (motif from Fig. 2e) to other PAS (statistical control) depending on how the individual PAS usage changes from qNSC to aNSCs in wildtype controls (left group) and APLP1-/- mice (right group), trend estimated by spline regression, ANOVA interaction test. **f**, Bcas2 (breast cancer amplified sequence 2) is shorter in APLP1-/-, left sub-panel, summed raw 3’ mapping positions, right sub-panels, spline regression of proximal to distal PAS usage over NSC lineage progression (pseudotime). **g**, FACS quantification of NSC numbers in APLP1-/- and wildtype mice (8-12 week adult mice), roughly 200 NSCs fewer in APLP1-/-, NSCs alive per mouse estimated by flow cytometry (FACS), surface markers (GLAST+, Prom+, Ter119-, CD45-, O4-), t-test, Welch’s correction, two-sided; error bars indicate standard deviation. **h**, Alterations in communication: APLP1-/- mouse pups, vocalizations recorded on postnatal days 5 (left) and 8 (right), classified by the *DeepSqueak* tool into different call types, average call numbers per genotype, color code: red = APLP1-/-, black = wildtype, red boxes highlight strongest differences. **i**, Affection and sociability in APLP1-/- mice, left sub-panel, time spent with a littermate, sociability is on average roughly 50% higher in APLP1-/-, right sub-panel, social novelty assayed in the three chamber test, novelty index indicates the time spent with an intruder mouse, roughly 50% lower in APLP1-/- mice, t-test, Welch’s correction, two-sided; error bars indicate standard deviation.

To explore a potential role of CPEB4 and other associated polyadenylation factors in APA regulation, we first examined expression of these factors in L2/3 excitatory neurons. In these neurons, CPEB4 and NUDT21 expression showed a tendency for upregulation in ASD patients compared to controls (Fig. 3k and Extended Data Fig. 4g). NUDT21 is a core regulator of APA, which was shown to be involved in neuronal fate determination of pluripotent stem cells^11^. Partial reduction of NUDT21 protein leads to altered APA in hundreds of genes and intellectual disability^30^. Surprisingly, we also found changes in expression of the synaptic adhesion molecule, amyloid beta precursor like protein 1 (APLP1). Interestingly, in neurons, APLP1 is used for synaptic compartmentalization of translation through its binding to CPEB1 that mediates poly(A)-tail lengthening^31^. Thus, we wondered if this factor would also be involved in regulation of APA by CPEB4. Interestingly, also in the mouse data set, Aplp1 showed similar regulation of expression as Cpeb4, namely, highest in quiescent NSCs and neuroblasts (Fig. 3l Extend Fig. 4h). Hence, to study potential alteration of APA if APLP1 is absent, we sequenced the transcriptomes of the olfactory bulb lineage from Aplp1-/- mice and their wild-type (WT) counterparts (Fig. 4a, Methods & Extended Data Fig. 5a). First, differential expression analysis between both genotypes revealed changes in genes associated with synapse organization, transmembrane transport, glial differentiation, axon development and behavior among others (Extended Data Fig. 5b & Supplementary Table. 2). Second, we examined regulation of 3’UTR length along the lineage. 3’UTR changes in these WT mice were in good agreement with observed changes in previously used WTs (R = 0.68, Extended Data Fig. 5c). We compared PAS selection between the two genotypes by multinomial regression and found over 900 genes showing differential PAS usage (Fig. 4b, Methods, ANODEV). In Fig. 4b the effect size (y-axis) is quantified for each gene by the earth mover’s distance (Extended Data Fig. 5d) between the 3’UTR length distributions (example genes in Extended Data Fig. 5e) of either genotype. We performed GSEA on the result of this regression, ranking genes by regression scores explained by genotype (see Methods for details), and found enrichment of genes localized to axons and dendrites (Fig. 4c, upper panel), and most importantly, of ASD risk genes (Fig. 4c, lower panel).

We next assessed whether genes exhibiting significantly differential PAS usage between Aplp1-/- and WT (as shown in Fig. 4b) would rather bind CPEB4 than CPEB1. To this end we use the list of CPEB1 and CPEB4 binders in neurons^29^. Approximately 50% of these genes that change their 3’UTR usage in Aplp1-/- vs. WT bind CPEB4 (Fig. 4d). This corresponds to a two-fold enrichment over the background of CPEB4 substrates, while CPEB1 substrates show no enrichment (Fig. 4d). In addition, we found a comparable enrichment in our NSC RIP assay, however the total number of CPEB4 substrates was here lower (Extended Data Fig. 5f). As in the previous data set, selection of PAS containing aCPE motif was favored in the transition from quiescent to active NSCs in WT controls. However, this bias was absent in the Aplp1-/- neuronal lineage (Fig. 4e). As an example, Fig. 4f depicts the gene Bcas2 (breast cancer amplified sequence 2), which was associated with ASD^32^. In the WT, its proximal PAS bearing three up-stream CPE motifs increases in the transition from qNSCs to aNSCs, while in APLP1-/- the distal PAS is poorly selected (Fig. 4f).

Next, we studied the consequences of the APLP1 deficiency at the cellular and behavioral level. First, we observed a lower density of qNSC transcriptomes and higher density of neuroblast transcriptomes in Aplp1-/- cells as compared to WT ones (Extended Data Fig. 5g). FACS quantification of the total number of NSCs isolated from the vSVZ revealed a decreased number of NSC in APLP1-/- compared to WT mice (Fig. 4g). Altogether, this suggests a higher rate of activation of NSC for generation of neurons would result in faster depletion of the NSC pool. Finally, we addressed if dysregulated APA along the neuronal lineage would affect the social behavior of mice, as it is the case in humans with ASD. To this end, we recorded the number of calls triggered by isolation from the mother at postnatal days 5 and 8 (P5 and P8), which is a measure of current affective state and motivation of pups. Calls were classified into different types. At P5, Aplp1-/- pups registered twice as many flat calls compared to WT pups of the same age (Fig. 4h). At P8, call types such as “upward ramp”- and “step up” showed reduced frequency in Aplp1-/- mice as compared to controls. At older ages, the three-chamber social interaction test revealed a decreased social novelty index in adult Aplp1-/- males than WT controls (Fig. 4i). In addition, we observed a tendency to higher anxiety in the plus maze and the open field test in APLP1-/- as compared to controls (Extended Data Fig. 5h). In summary, Aplp1-/- mice exhibit changes in the affective state and sociability that are commonly used as surrogate symptoms for autistic or neurodevelopmental disorders in humans^33–35^.

To summarize, we carried out a functional gene study with the aim of describing 3’UTR regulation in the context of the NSC lineage progression and in disease. We indeed demonstrate regulation of APA along the neuronal lineage in genes associated with developmental disorders such as ASD. Further, we observed an overall lengthening of 3’UTR in transcripts of neurons from ASD patients as compared to healthy individuals. Regulated 3’UTRs exhibited an enrichment of CPE motifs that bind to CPEB4. In NSCs shortening 3’UTRs results in increased protein outcome. Finally, studies of Aplp1-/- mice suggest that CPEB4 works together with APLP1 to control APA along the neurogenic lineage. Absence of APLP1 decreases selection of CPE-PAS that potentially contributes to dysregulation of posttranscriptional regulation of gene expression in Aplp1-/- neuronal lineage and autistic-like traits in mice as compared to their WT counterparts. Our study paves the way for studies further addressing the upstream regulators of APA in neurodevelopment potentially dysregulated in neurodevelopmental disorders.

## ACKNOWLEDGEMENTS

We thank U. Müller and the members of her lab for providing us with Aplp1-/- mice, the TAC members M. Boutros, M. E. Al Shukairi for technical assistance and the members of the Martin-Villalba laboratory for critical comments. This work was supported by the German Cancer Research Center (DKFZ), the University of Heidelberg and the DFG (SFB873).

## AUTHOR CONTRIBUTION

Project concept, design, and oversight A.M.-V, S.A.; M.G. conducted computational analysis supervised by S.A., W.H., A.M.-V.; N.G.-O conducted studies of APLP1 mutant and wt mice; CPEB4 expression analysis, APA validation studies; A.D.M, A.B, R.P, conducted studies of CPEB4 NSCs supervised by R.M. and I.F.; D.F. and J.K. conducted proteomics analysis; U.M. generated and provided APLP1 mutants; Writing-Review & Editing, A.M.V., S.A., W.H, M.G, N.G.-O.

## AUTHOR INFORMATION

Raw sequence data is available from GEO (Accession number: GSE). Complete documented software is available as a supplementary file. The complete R/Bioconductor code used for the analysis will be made available upon acceptance on the authors’ webpage (https://martin-villalba-lab.github.io/). The authors declare no competing financial interests. Correspondence and requests for materials should be addressed to AMV.

## METHODS

### Mouse strains

The WT mice used in this study were C57BL/6 male mice. For experiments involving the role of CPEB4 and APLP1, knockout mice for the respective genes were used along with their corresponding WT mice as controls. The mice used were 8-12 weeks of age. The mice were subjected to a 12h light/dark cycle and were housed in the animal facility at the German Cancer Research Center (DKFZ) with free access to food and water. All experiments conducted were in accordance with the guidelines prescribed and approved by the *Regierungspräsidium Karlsruhe*, Germany.

### Isolation and culture of NSCs *in vitro*

Single cell suspensions were prepared from microdissected vSVZs using the Neural Tissue Dissociation Kit with Papain (Miltenyi). The vSVZs from each mouse (3 mice/biological replicates) were processed separately. Isolated NSCs were cultured in proliferating media; Neurobasal-A medium (NBM-A) containing B27 supplement (2%, GIBCO), L-Glutamine (1%), heparin (2 μg/ml), human FGF (20 ng/μl, ReliaTech) and human EGF (Promokine). Cells were not used beyond 10 passages.

### Induction of quiescence *in vitro* with BMP4

Labtek chamber slides/culture dishes were coated with Poly D-Lysine (1 μg/ml, overnight) and Laminin (10 μg/ml, 2 hrs). The *in vitro* NSCs were then seeded onto pre-coated plates (30,000 cells/cm^2^) in proliferating media. On the following day the media was replaced with quiescence media in which EGF was exchanged for BMP4 (50 ng/ml, R&D Systems). The cells were cultured for a total of 3 days with fresh media exchanged every 2 days. A lower seeding density (10,000 cells/cm^2^) of active NSCs was used in order to keep the final cell numbers comparable to the BMP4 treated quiescent NSCs and were cultured in proliferating medium^36^.

### Immunofluorescence

Cells cultured in chamber slides coated with PDL & Laminin were fixed in 4% PFA for 20 mins. Following fixation, the cells were blocked in 0.3% Triton X-100, 3% horse serum for 45 min at RT to avoid any unspecific staining. The cells were then incubated with the primary antibody, anti-Ki67 (rabbit, novus, 1:200), overnight at 4°C. A secondary antibody (Donkey anti-rabbit 546; 1:1000) was used.

### Microscopy

A Leica TCS SP5 microscope with a UV diode (405 nm) and a helium-neon 561nm laser was used to acquire confocal images. For the immunofluorescence imaging of BMP4 induced quiescent vs. EGF control in vitro cells 6-12 z-stacks (1μm) were obtained at a resolution of 1024×1024, 2-3 images per treatment type/biological replicate were imaged and the results averages across all images. The images were captured using a 40x oil-immersed objective. The number of Ki67 positive and negative cells were quantified using the imageJ plug-in Cell Counter on a max projection image of the different stacks obtained.

### Preparing cells for 10x Chromium

For *in vitro* cells, after treatment with BPM4/EGF for 3 days, the cells were trypsinized to prepare a single cell suspension. The 3 biological replicates per treatment were combined in equal numbers to achieve a cell density of 200 cells/μl. For the Aplp1-/- vs. WT, single-cells suspensions were prepared as described above from vSVZs. Each mouse was processed separately. We FACS sorted CD45^**-**^ O4^**-**^ Ter119^**-**^ GLAST^**+**^ cells to enrich for the neural stem cell lineage. In order to keep the replicate information from individual mice for the Aplp1-/- vs. WT mice, tissue from each mouse was processed separately with the MACS Neural Tissue Dissociation kit (T). Each of these samples were stained with O4 APC-vio770, Ter199 APC-Cy7, CD-45 APC-Cy7, GLAST-PE. Unique cell hashing antibody was added to each sample (BioLegend) in order to preserve the replicate identity. The samples after sorting were then subjected to the 10x protocol (v2) to generate the GEMs and were further processed to generate 10x libraries. These libraries were sequenced on the NovaSeq 6000 PE 150 (for the 3’UTR analysis).

### FACS analysis of APLP-/- and WT mice

In order to quantify the NSC numbers in APLP1-/- and WT mice (8-13 week old adults), single cells suspensions were prepared as described above from vSVZs. Each mouse was processed separately and the single cell suspensions were stained for O4 APC-vio770, Ter-199 APC-Cy7, CD-45 APC-Cy7, GLAST-PE, Prom1-APC, and dead cell exclusion by Sytox Blue. NSCs were defined as Sytox^**-**^ CD45^**-**^ O4^**-**^ Ter119^**-**^ GLAST^**+**^ Prom1^**+**^ cells. Samples were run on FACS Canto, and FACS Diva software was used for analysis of the data.

### Western Blot

Total protein lysates were prepared from BMP4 and EGF treated NSCs. 500,000 cells per sample were used to prepare the lysates. For CPEB4 localization cytoplasmic and nuclear protein fractions were extracted from BMP4 and EGF treated NSCs (around 2 million cells per sample) using NE-PER™ Nuclear and Cytoplasmic Extraction Reagents (Thermo Scientific™). GAPDH and Histone H3 were used as markers for cytoplasmic and nuclear fractions respectively. The following primary antibodies were used for the western blot: anti-CPEB4 (149c) (Mouse, 1:500); Anti-Vinculin (Abcam, ab129002, Rabbit, 1:5000), GAPDH (Cell Signaling, Rabbit, 1:1000), Histone H3 (Cell Signaling, Rabbit, 1:1000)

### Behavioral Experiments

All behavioral experiments were conducted at the Interdisciplinary Neurobehavioral Core (INBC) of Heidelberg University.

### Ultrasonic Vocalization (USV)

USVs were recorded as described previously^37^. Infant mice at two different developmental stages (postnatal day, PD 5 and 8) were separated from the mother and their USVs were recorded. The pups were placed in an empty glass container (6 × 9.5 × 7.5 cm) filled at the bottom with bedding material. This container was then placed inside a white acryl open field box (40 × 40 × 40 cm) and the roof was covered with a Plexiglas lid to dampen ambient noise. A microphone was placed 30 cm above the bedding through the roof to record the USVs. The USVs were recorded for duration of 5 min and the pups were returned back to their cage. Care was taken to always start the experiment at the same time of the day and the same pups were used to record USVs for both P5 and P8. The number of pups used in this study was 15 WTr6 (11 male and 4 females) and 17 APLP1-/- (11 males and 6 females).

### USV Recording and Analysis

The recordings were conducted with equipment and software provided by Avisoft Bioacoustics (Berlin, Germany). Ultrasonic condenser microphones CM16/CMPQ were placed 30 cm above the floor of the testing area. The acoustic signals were recorded, amplified and digitized at 250 kHz with 16-bit resolution by the Ultrasound Gate 416 Hz USB audio device. The USVs were analyzed and classified using the DeepSqueak^38^ (MATLAB package. The call classification was done using DeepSqueak supervised classification of call types.

### Social Interaction test in adolescent mice

Social interaction between littermates was assessed as described before^39^ in adolescent mice (PD 21) between the same sex and litter. One mouse from each gender was isolated for 24 h from their littermates. The test included two trials. The habituation trial involved each test mouse being placed in the arena (open field box; 40 × 40 × 40 cm) and allowed to habituate for 2 min. In the sociability trial the isolated mouse was reunited with each of the testing mice and their interactions were tracked by EthoVision XT video tracking software for a duration of 5 min. USVs were recorded in both trails. Proximity of interaction was defined as a distance between two mice that was less than 10 cm. Cumulative duration of proximity was calculated for each pair of mice. The number of mice used in this study was 23 WTr6 (16 male and 7 females) and 15 APLP1-/- (10 males and 5 females).

### Three-Chamber Sociability and Social Novelty Test

Sociability and social novelty for Aplp1-/- vs. WTr6 mice were tested in young mice (Post natal day 40) as described previously^33,34,40^. Interactions were restricted to mice of the same gender and litter. The experiment was conducted in a clear Plexiglas rectangular box that is divided into three chambers with small openings connecting them. In the left and right chambers a cylindrical caged enclosure was placed in the center. The experiment was conducted in 3 trials lasting 5 min each and the testing mouse was placed back in its cage after every trial. An interval of 15 min was given between trials so as not to stress the mice. In the habituation trail the testing mouse was allowed to explore the arena for the duration of the trial. In the sociability trial a littermate isolated 24hrs prior was placed in the caged enclosure in the right chamber. The testing mouse was then introduced back to the central one and its interactions with its littermate were monitored. The final trial was to check for social novelty where an intruder was introduced in the caged enclosure in the left chamber. The test mouse was then introduced back to the central chamber and its interactions with both its littermate as well as the novel intruder mouse are monitored. Sygnis Tracker software was used to track the movements of each mouse in the arena. The time spent in the vicinity of an empty cup, littermate or intruder were used to estimate a sociability and social novelty index. The number of mice used in this study was 24 WTr6 (15 male and 9 females) and 16 APLP1-/- (10 males and 6 females).

### Proteome sample preparation

Cell pellets were lysed in 20 μl lysis buffer composed of 50 mM ammonium bicarbonate (AmBic), 1% sodium dodecyl sulfate (SDS), 10 mM tris (2-carboxyethyl)phosphine (TCEP), 40 mM chloroacetamide (CAA) and protease inhibitor cocktail without EDTA (Roche). Pellets were suspended using a vortex mixer; samples were frozen in liquid nitrogen, thawed and subjected to sonication on melting ice using a probe sonicator (Branson) with 10% output for 10 seconds. A single-step reduction and alkylation was carried out at 95 °C for 5 minutes. Samples were subsequently cooled to room temperature and briefly centrifuged. Proteome samples were prepared according to the SP3 protocol^41,42^ with modifications. Briefly, GE Healthcare Sera-Mag Speed Beads A and B (Thermo Fisher Scientific, Germany) were equilibrated to room temperature. Magnetic beads were prepared for use by combining 20 μl of beads A and B with 160 μl of ddH2O. Beads were settled on a magnetic rack for 2 minutes and the supernatant was discarded. Beads were washed twice more with 200 μl ddH2O and resuspended in 20 μL ddH2O. For protein purification, 2 μl of prepared beads and 22 μl of 100% acetonitrile (ACN) were added to each sample. Protein binding to the beads continued for 18 minutes, followed by immobilizing the beads for 2 minutes on a magnetic rack. The supernatant was discarded, beads were washed twice with 200 μl of 70% ethanol (EtOH), once with 180 μl of 100% ACN and air-dried without displacing them from the magnetic rack. Beads were carefully reconstituted in 10 µl of 100 mM AmBic and sonicated for 5 minutes in a water bath. Protein digestion was performed overnight at 37 °C in a shaking incubator using 0.12 μg trypsin (Promega) at an enzyme to protein ratio of 1:50 (protein content was calculated using independent samples). For peptide purification, 210ul of 100% ACN was added to each digested sample. Peptide binding to the beads continued for 18 minutes, followed by immobilizing the beads for 2 minutes on a magnetic rack. The supernatant was discarded, beads were washed twice with 180 μl of 100% ACN and air-dried without displacing them from the magnetic rack. To recover peptides, beads were suspended in 10 μl of 0.1% formic acid (FA) by water bath sonication for 5 minutes. Samples were briefly centrifuged and placed on a magnetic rack. Supernatants were transferred to new tubes and settled on a magnetic rack again to ensure a complete removal of beads before further processing of peptide-enriched supernatants.

### LC-MS/MS

Peptide samples were analysed with a Orbitrap Fusion mass spectrophotometers connected to EASY-nLC 1200 liquid chromatography system via a nanoelectrospray ion source (Thermo Fisher Scientific). Peptides were separated with a C18 (75 µmx 250 mm, 1.7 µm) UPLC column (Thermo Fisher Scientific) using a stepwise 105-min gradient, from 4% to 100% (v/v) acetonitrile in 0.1% (v/v) formic acid at a flow rate of 300 nl/min. Orbitrap Fusion was operated using Orbitrap as MS1 analyser and Ion Trap as MS2 analyser. Charge states of two, three or four were allowed for fragmentation. Essential MS settings were: full MS: AGC = 10E6, maximum ion time = 50 ms, m/z range = 375-1500, resolution = 60 000 FWHM; MS2: AGC = 1E4, maximum ion time = 50 ms, minimum signal threshold = 1000, dynamic exclusion time = 40 s, isolation width = 2 m/z (1.6 m/z) and normalized collision energy = 33%.

### Protein identification and quantification

The raw mass spectrometry data were processed with MaxQuant^43^ (version 1.6.0.16). The MS/MS spectra data were searched with the Andromeda search engine against the mouse canonical reviewed proteins of the Uniprot database (version 2018_02). A minimum of one peptide was required for protein identification. The false discovery rate (FDR) at peptide and protein level was set to 0.01. The enzyme specificity was set to trypsin/P and 2 missed cleavages were allowed. Cysteine carbamidomethylation was set as a fixed modification, while protein N-terminal acetylation and methionine oxidation were included as variable modifications. The minimal peptide length was set to 7 amino acids. For the LC-MS/MS a maximal allowed tolerance was set to FTMS 20 ppm and ion trap MS (ITMS) 0.5 Da. Protein quantification was based on the MaxLFQ label-free quantification algorithm^44^.

Perseus^45^ (version 1.5.3.0) was used for differential analysis of protein expression. Potential contaminants and reverse database hits were filtered out. MaxLFQ values were log2 transformed and corrected by the median log2 value per sample. Proteins quantified in 2 out of 3 biological replicates in each condition were subjected to differential expression analysis by means of two-sample t-test with permutation-based FDR multiple test correction. Adjusted p-values equal or lower than 0.05 pointed to statistically significant protein expression change between BMP4 treated and control neuronal stem cells. As a measure for posttranscriptional regulation, matched genes between proteome and transcriptome were combined into a single value referred as translation index (illustrated in Extended Data Fig. 3f). For each condition (EGF, BMP4), LFQ values were divided by summed UMIs and subsequently logarithmized and centered around 0.

### Statistical analysis

Differences between groups measured as correlation coefficients or fold-changes were assessed using non-parametric Wilcoxon rank sum tests (in *R*). T-tests were conducted when appropriate (R, GraphPad Prism), Welch’s correction and pairwise t-test for multiple groups (in *R*). Multinomial test were computed in *R* using the *nnet* package and related ANOVA functions. Correlation coefficients and tests were estimated utilizing Pearson’s method (in *R*). Statistics that were computed on one sample (i.e. on single cells) and not biological replicates were reported as nominal p-values.

### Mapping of 3’UTRs and statistical analysis of APA

For the analysis of 3’UTRs, 10X reads (100bp or 150bp length) were mapped in paired-end mode using bowtie2^46^ (version 2.3.5.1) with the option *--very-sensitive-local* to the mm10 genome. Subsequently, alignments mapping to regions annotated as 3’UTRs from the Ensembl built were extracted. Only primary and alignments mapped as proper pairs were considered for further analysis. Per region, the most 3’ mapping position of read 1 was translated from *CIGAR* strings to local coordinates in transcripts using custom scripts. To correct for outliers, the median mapping position was counted for duplicated reads using the unique molecular identifier (UMI) information. This workflow was executed in the R environment by loading the *BAM* files with the R packages *GenomicAlignments and GenomicFeatures*^47^. Results were stored in a region-ordered list. For each entry in the list, 3’tail peaks were annotated in an unsupervised manner by clustering the non-zero positions, *i*.*e*. UMI counts of 3’UTR ends, using the functions *hclust()* and *cutree()*. A 3’tail peak was called if it was supported by at least 3 UMIs and had a distance to other peaks of at least 50 bp considering the mean peak width of around 20 bp. The outcome was a gene ordered list storing count tables with respect to 3’ tail peaks and single cells as well as other features as S4 objects. As a validation, the distance of peak centers to the closest canonical poly(A) signal (AAUAAA) in the 3’UTR reference sequence was computed.

Subsequently, multinomial regression on each gene that featured two or more detected peaks (APA) was computed utilizing the *multinom()* function from the R package *nnet* .Explicitly, the logit function for this regression approach is given as:

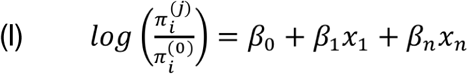

Where *π* in (I) are the probabilities for *j* categories (annotated 3’peaks), the superscript *0* the reference category (first and therefore most proximal peak in *nnet*, R package) and *x* the response. UMIs were used as weights for each observation (in *x*). Parameters *β* were determined by fitting the model. In particular, the changes in local odds were modeled as basis splines (B-splines over *x*) of the predictor variable with 3 degrees of freedom. Gene-wise, ANOVA tests were executed on a full model including the pseudo-time as predictor and a reduced model with a constant predictor variable. For every APA gene, the likelihood ratio (LR) statistics were extracted and utilized to score the genes. This was done to test whether APA depends on pseudo-time representing the NSC lineage progression. For visualization and estimation of the dynamics of 3’UTR usage, the fitted beta-parameters (*β* from I) were used for back-transformation of B-splines (R package *splines*) to get smooth curves per 3’peak estimating the fraction of UMIs of the given peak over pseudotime.

**Analysis of 3’UTR changes for APLP1-/- vs. WT**

Mapping of 3’UTRs was carried out as described in the previous paragraph. As measure for the difference in 3’UTR usage between APLP1-/- to WT the Earth Mover’s distance (EMD) was computed (see also Extended Data Fig. 5d). For every APA gene a matrix (illustrated in Fig. 4a) was defined. The EMD was computed as:

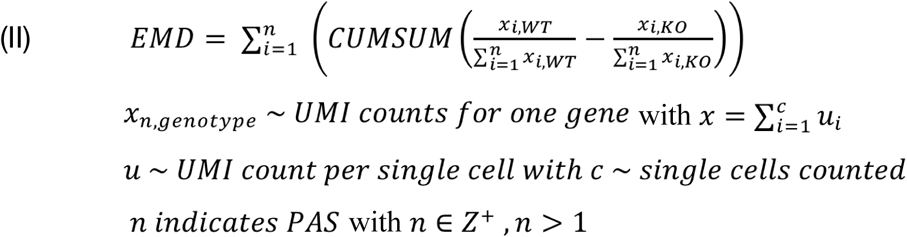

Here, EMD < 0 indicates shortening of 3’UTRs in APLP1-/- compared to WT, EMD > 0 lengthening. Statistical significance was tested using a log-likelihood test with a full model including the predictor “genotype” vs. a reduced one with a constant predictor (*multinom* function from *nnet*). The 3’peak counts per biological replicate were considered as statistical unit. Since this is a multinomial test (not an ordinal), also changes in intermediate 3’peaks are detected. For simplification and reporting, a gene was called longer or shorter in the genotype comparison if FDR < 5% and EMD consistently higher or lower then 0 across biological replicates.

### Enrichment Analysis of Gene Sets and Motifs

Enrichment analysis employed the R packages *ClusterProfiler*^48^, *DOSE* and *ReactomePA*. Gene set enrichment analysis (GSEA) was used to determine whether annotated gene sets (categories from Gene Ontology (GO), Disease Ontology (DO)^49^, Disease Gene Network (DGN)^50^ or the Reactome Pathway database) were over-represented at the top or bottom of a ranked gene list. Significance was computed by permutation tests according to the GSEA method with p-value adjustment for multiple testing (*i*.*e*. multiple categories). A normalized enrichment score (NES) was reported since this value includes the mean of all permutations and accounts for correlations to other gene sets. Input gene lists were ranked either by the multinomial regression statistics (evidence for 3’UTR changes), log2-fold-changes (CPEB4 immunoprecipitation and gene expression in APLP1-/- vs. WT) as well as 3’UTR lengthening trends. For DO and DGN, mouse gene identifiers were translated to human orthologues using the *getLDS* function from *biomRt*. For the translation, duplicated gene identifiers were removed, non-translatable ones not considered and all multi-match identifiers were used.

Two gene categories were of special interest comprising the list of 83 genes related to APA regulation with the associated molecular functions from GO: “mRNA polyadenylation” (GO:0006378) and/or “mRNA 3’end processing” (GO:0031124).

Motif analysis in 3’UTRs was performed applying the tool homer2^20^ (v4.9) for de-novo motif detection with the command: *homer2 denovo -len 7 -i fgd*.*fasta -b bck*.*fasta*. As background sequence contexts of 3’peaks from genes with only one 3’peak (constant 3’UTR) were used. As foreground, 3’peaks from APA that show clear variation in the UMI fraction with pseudotime were considered, irrespective whether they increase or decrease. To describe the effect of CPEB4 binding on 3’UTR length choices, a CPE motif (as regular expression: *TTTTGT*|*TTTTGAT*|*TTTTAGT*) was used If the 3’peak had at least one occurrence of this motif in a distance of 1 to 50 bp upstream (in its 3’UTR sequence) it was classified as CPE containing 3’peak.

### Analysis of pseudotime and expression levels in NSCs

Pseudotime was computed using the standard workflow from the R package *monocle* 2^51^ and according to the method described in Kalamakis *et al*. 2019. Features were selected by choosing genes with the highest dispersion (variance) in the respective dataset. Importantly, pseudotime was computed separately three times for each sequencing experiment: *in-vivo* NSCs (young & old), *in-vitro* NSCs (EGF & BMP4) and APLP1 (WT & APLP1-/-).

For the expression level comparison of *in-vivo* to *in-vitro* NSCs, single cells from both systems were matched by subsampling *in-vitro* NSCs. This way, every *in-vivo* cell has an *in-vitro* counterpart in pseudo-time. For the gene-wise correlation tests (Pearson’s moment correlation, one-sided), the expression of genes between pseudotime ordered cells (*in-vivo* vs. *in-vitro*) were computed. For the UMAP embedding the method from Haghverdi 2018 for single cell batch correction was applied (MNN - mutual nearest neighbors). Distances of single cells in expression space were visualized with the Sleepwalk^52^ tool as Euclidean distance.

### Preparation of CPEB4 RNA immunoprecipitation (RIP)

For RIP-seq analysis, adult subventricular NSC cultures were obtained from 2-months-old CPEB4-/- and WT mice (n = 2 per genotype) and sub-cultured as previously described (Belenguer *et al*., 2016). Secondary neurospheres grown for 3 days (3-4 million cells) were harvested and crosslinked with 0.5% formaldehyde (Thermo Fisher, cat. no. 28908) in 0.1 M DPBS for 5 min at RT with soft agitation. Crosslinking was stopped with 0.25 M glycine for 4 min and washed twice with ice-cold DPBS. Pellets were processed as previously described (Maillo et al., 2017). Briefly, pellets were lysed with RIPA buffer (plus protease and RNase inhibitors) and then sonicated. After centrifugation, supernatants were collected, precleared, and immunoprecipitated with 10 μg of anti-CPEB4 antibody (Abcam, ab83009), or rabbit IgG (Sigma) bound to 50 μl of Dynabeads Protein A (Invitrogen). After treatment with Proteinase K, RNA was extracted by standard phenol–chloroform protocol. Samples were processed at IRB Functional Genomics Facility following standard procedures.

### Analysis of CPEB4 RNA immunoprecipitation results

For CPEB4-RIP, reads were mapped to the mm10 genome (single reads, 50 bp) and counted from BAM files using the recommended Bioconductor workflow (function *SummarizeOverlaps* with mode *‘Union’*). Duplicated reads were removed. Subsequently, DESeq2 was applied to compute the log2-fold-changes between the IP fraction and controls. As validation for the CPEB4-specific CPE motif: for every 3’UTR the read coverage was fitted as generalized linear model (GLM) with the “poisson” family distribution as follows: *IP ∼ IgG + genotype* (controls: IgG, Immune Globulin and CPEB4-/- IP fraction). The residuals from the model fits were considered and averaged over genes relative to the CPE motif or a random position/motif as sanity check. (Positive residuals indicate that the WT IP is higher than the control fractions.)

### Assignment of biological replicates (cell-hashing)

The abundance of hash-tags with the sequences: *GGTCGAGAGCATTCA, CTTGCCGCATGTCAT, AAAGCATTCTTCACG* in APLP1-/- sequencing were analyzed using a custom R script employing the *ShortRead* package. At the starts of read 2 hash-tag sequences were matched allowing 1 mismatch. The single cell barcodes from read 1 were compared to those from the 10X genomics output. Each cell was assigned to an individual mouse (biological replicate) in case the cell had an over-representation of one hash-tag that was higher than 1.5-fold of the median hash-tag count that was detected within one cell.

### Analysis of single cell sequencing data from ASD patients

FASTQ-files were downloaded from GEO (accession ID: PRJNA434002) and subsequently processed with the 10X genomics pipeline (*cellranger count*, version 2.2.0). Reads were mapped against the human genome hg38 (GRCh38). UMI counts per gene were summed over single cells grouped by cell-type (cell-type annotation adopted from the original paper) and individual. Alterations in gene expression were assessed applying DESeq2 (model formula: ∼ brain-region + sex + diagnosis). Differential expression was computed separately for every cell-type with the standard multiple testing correction from DESeq2.

For every 3’UTR region in the human genome (gene annotation from the R package: *TxDb*.*Hsapiens*.*UCSC*.*hg38*.*knownGene*) the most 3’ mapping position of read 2 was counted (similar as the workflow for mouse data using a custom script). 3’UTR lengths were averaged for each individual and cell-type over single cell, respectively. ANOVA methods for linear models (in R) were applied on these values, thus computing p-values based on samples (unit of replication) and not on single cells. In detail, it was tested whether the inclusion of the predictor “diagnosis” (full model) had a significant effect on the model parameters and therefore the distribution of 3’mapping positions. Explicitly, the model was formulated for every gene as for the differential expression test: *Length ∼ brain-region + sex + diagnosis*.

