## Extended Data Figures and Supplementary Data for "Single cell 3’UTR analysis identifies changes in alternative polyadenylation throughout neuronal differentiation and in autism"

##### *Extended Data Figure 1*

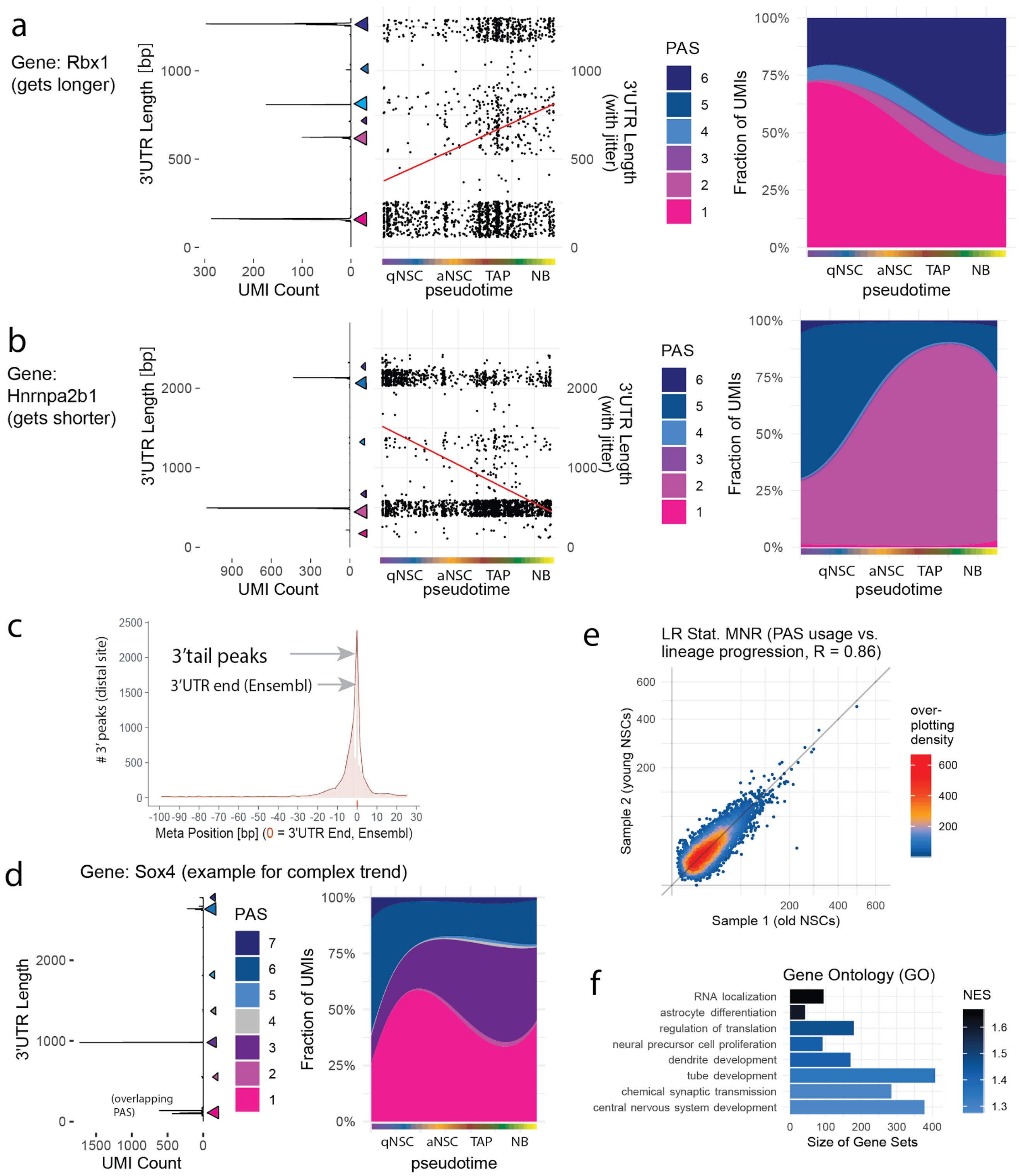

***Extended Data Figure 1*** **- Further exploration of 3'UTR changes along the NSC lineage.**

**a-b**, left sub-panels, raw 3' mapping pos. summed over cells, mid sub-panels, mean mapping pos. per cell (ordered in pseudotime as in Fig. 1a), linear regression lines fitted (in red), right sub-panels, approximation of changes in 3'UTR usage over pseudotime utilizing multinomial regression splines, color code as dark blue to red for most distal to most proximal PAS (color code corresponds to left sub-panels), **a**, Rbx1 (ring-box 1, also known as ROC1) example for 3'UTR lengthening, **b**, Hnrnpa2b1 (heterogeneous nuclear ribonucleoprotein A2/B1) example for 3'UTR shortening. **c**, Relative pos. of most distal 3' peak centers annotated with the applied peak calling algorithm (Methods) to 3'UTR ends from the Ensemble database (at pos. 0), shown as meta-gene analysis *i.e.* summed for expressed genes, pos. < 0 upstream of 3'UTR ends, positive pos. > 0 downstream. **d**, gene Sox4 left sub-panel, raw 3' mapping pos. summed over cells, right sub-panel, approximation of changes in 3'UTR usage over pseudotime utilizing multinomial regression splines, color code as dark blue to red for most distal to most proximal PAS (color code corresponds to left sub-panel). **e**, Log-likelihood statistic (LR stat.) for multinomial regression (Methods) independently computed from both representations of the NSC lineage ("sample 1" and "sample 2"), high if 3'UTR usage changes with lineage progression. **f**, Gene set enrichment analysis, Gene Ontology, ranked by LR stat. from **e**, a high normalized enrichment score (NES) associates respective gene categories with 3'UTR changes along the NSC lineage.

***Extended Data Figure 2***

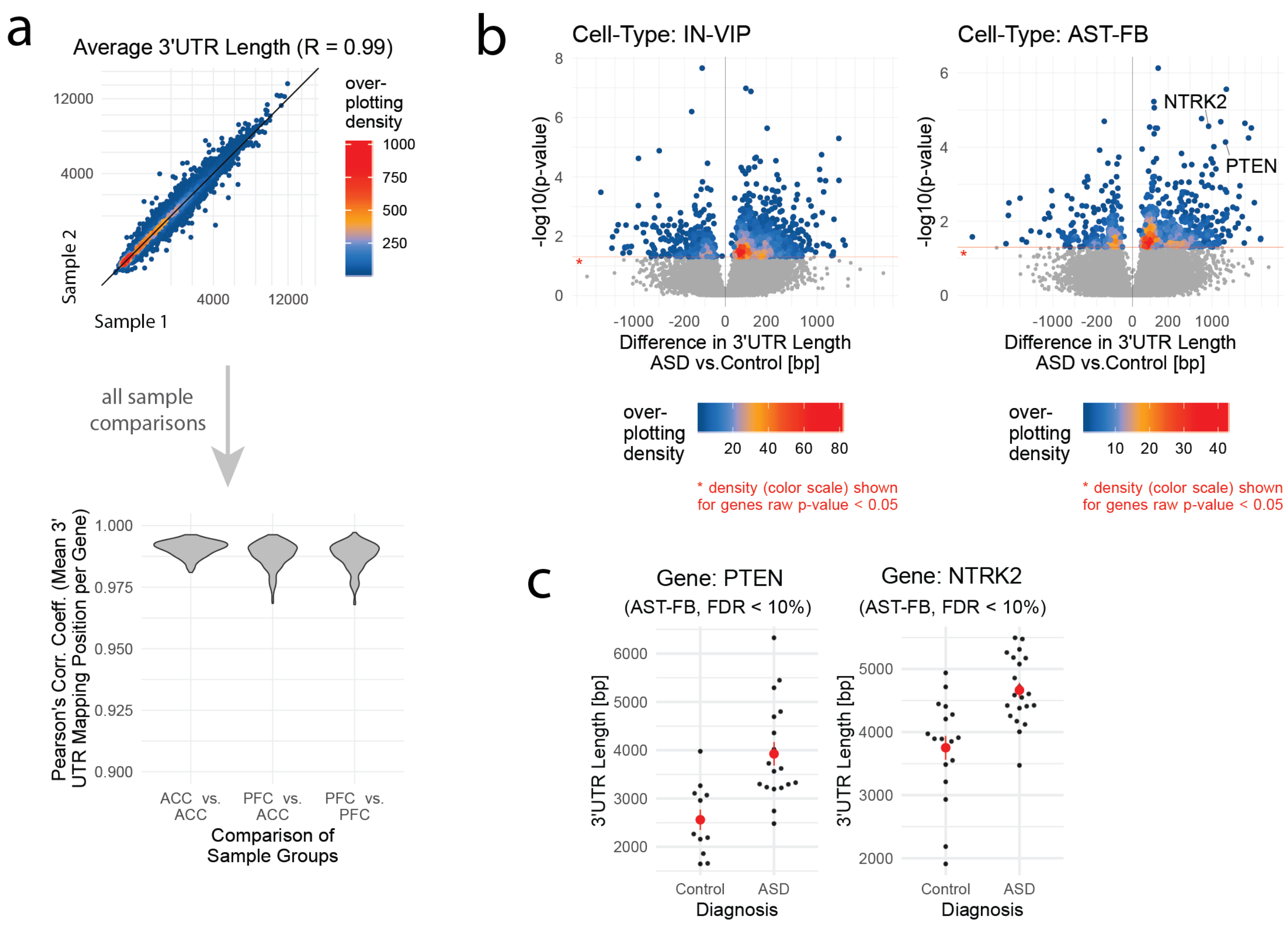

***Extended Data Figure 2*** **- Analysis of 3'UTR alterations in autism patients.**

**a,** Sample correlation scores, upper panel, representative scatterplot of mean 3'UTR mapping positions (across all single cells) per sample, color code to show density, lower panel, sample correlation computed for all possible sample combinations (shown as violin plots), groups: comparison of prefrontal cortex (PFC) and anterior cingulate cortex (ACC) samples. **b,** Volcano plots, 3'UTR alterations in interneurons (IN-VIP, left sub-panel) and astrocytes (AST-FB, right sub-panel) comparing ASD to healthy controls per gene, fold-chance in 3'UTR length (x-axis), p-value estimated from linear models (from Fig. 4c, y-axis), over-plotting color scale applied to genes with uncorrected p-values < 0.05 to show the overall trend. **c**, Example genes from b (AST-FB), every point average 3'UTR length over single cells per sample, red points group means, linear model (ANODEV) from Fig. 2a, left, PTEN (Phosphatase and tensin homolog), right, NTRK2 (Neurotrophic Receptor Tyrosine Kinase 2, also known as TRKB), both genes get longer in ASD vs. healthy control.

*Extended Data Figure 3*

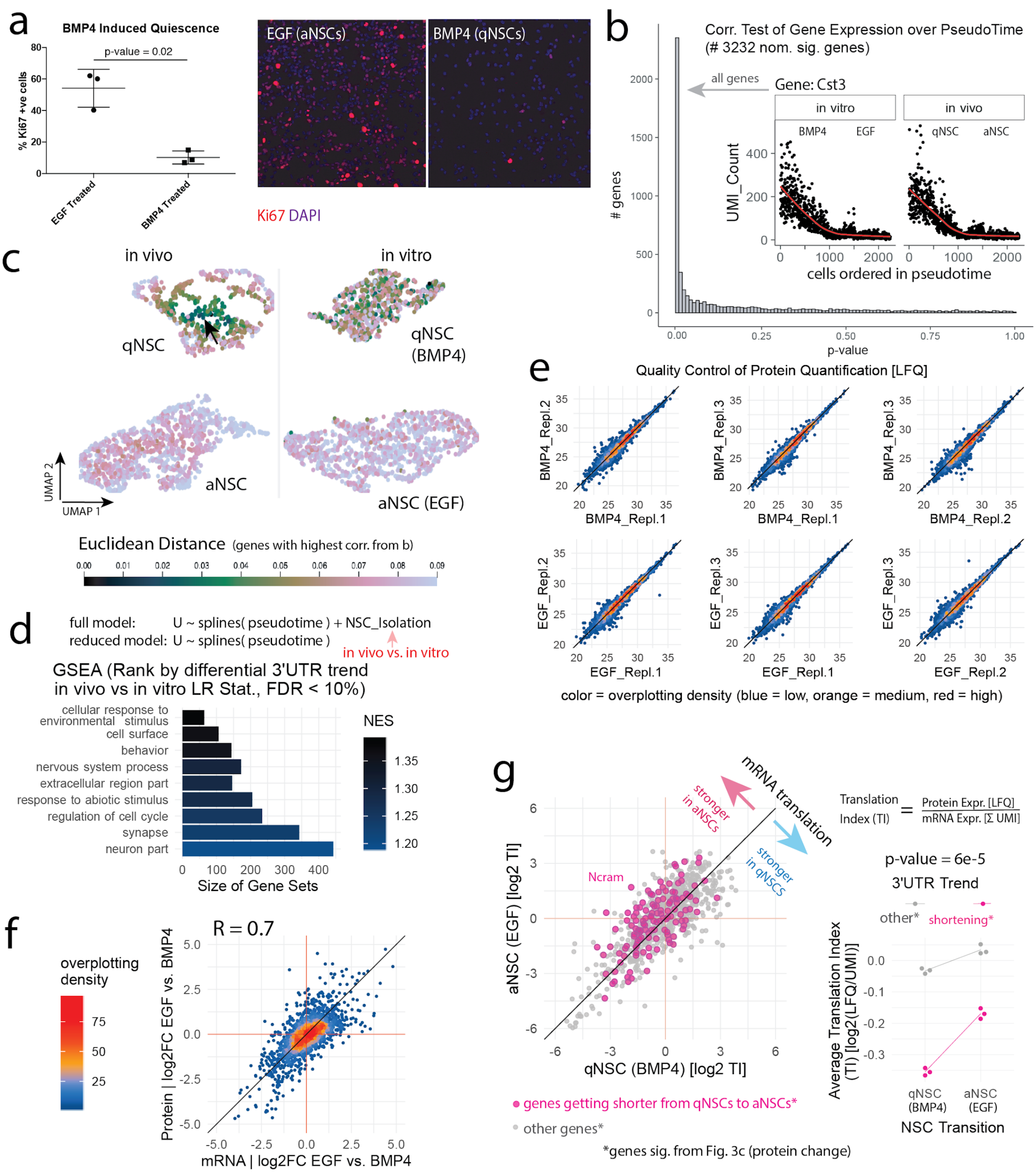

***Extended Data Figure 3* - Comparison of gene expression in *in vivo* vs. *in vitro* NSCs and quantification of protein amount in *in vitro* NSCs**

**a,** Staining for proliferation marker Ki67 (also known as Mki67) in cultured NSCs, treatment EGF (aNSCs) and BMP4 (qNSCs), left sub-panel, quantification of Ki67-postive cells (n = 3); error bars indicate standard deviation, right sub-panels, *in situ* staining example images with markers DAPI and Ki67. **b,** Statistical test whether genes follow the same expression trend along the NSC lineage in *in vitro* NSCs as they do *in vivo* NSCs, outer panel, p-value histogram, inner panel, example gene Cst3 (Cystatin C), every point is a single cell, red regression line to show the trend, nominal p-value. **c,** 2-D projection of *in vivo* and *in vitro* NSCs on the same UMAP, plot split for visualization, batch correction by mutual nearest neighbors (MNN), color code: Euclidian distance of *in vivo* qNSCs (see cursor position) to all other cells (black/ blue if cells are similar) by the sleepwalk tool. **d,** Gene set enrichment analysis, Gene Ontology, ranked by LR statistics testing for differential 3'UTR trend *in vivo* vs. *in vitro*, a high normalized enrichment score (NES) associates respective gene categories to show a different trend comparing *in vivo* to *in vitro*. **e**, Scatterplot matrix of biological replicates for protein quantification in cultured NSCs by mass spectrometry, upper row BMP4 (qNSCs), lower row EGF (aNSCs), quantified as LFQ values. **f**, Log fold-change, per gene: change in mRNAs levels (x-axis) against change in protein (y-axis) between aNSCs (EGF) and qNSCs (BMP4). **g**, left sub-panel, per gene translation index: non-degraded protein [LFQ] per non-degraded mRNA [UMIs], for aNSCs and qNSCs, shown for proteins significant from Fig. 3c, shortening genes in magenta, right sub-panel, average TI per gene cluster (UTR shortening vs. others), every point one biological replicate (from e), ANOVA interaction test.

***Extended Data Figure 4***

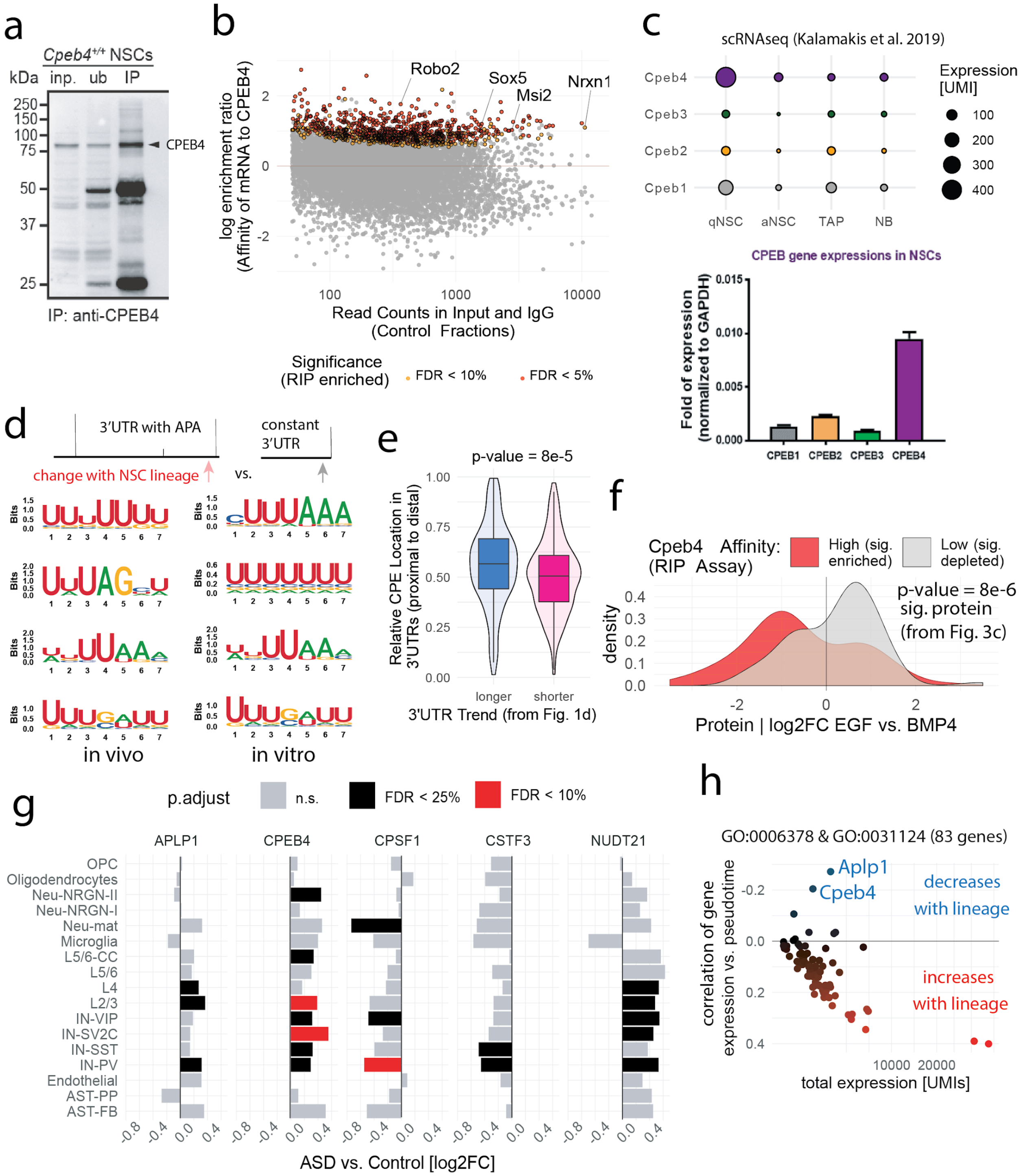

i

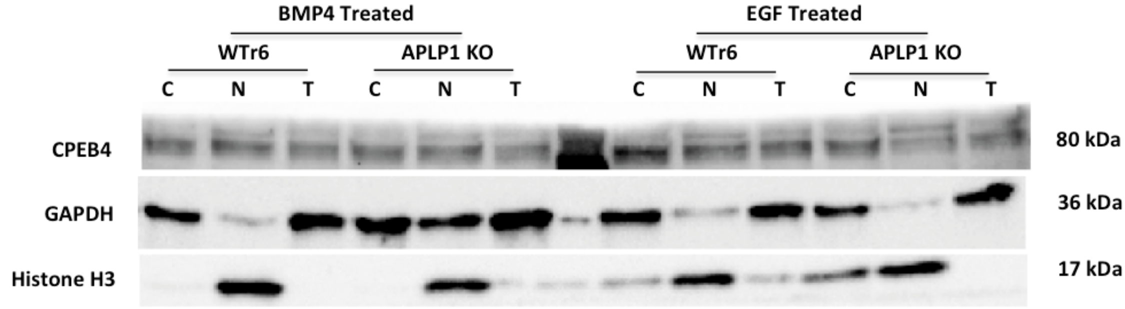

***Extended Data Figure 4*** **– CPEB4, RIP assay, protein expression, 3'UTR alterations and co-expression of with APLP1.**

**a,** Validation of CPEB4 RIP assay showing enrichment of CPEB4 in the IP fraction. **b,** Detection of CPEB4 substrates (CPEB4 binds mRNAs) by RNA immunoprecipitation (RIP), y-axis log-enrichment in CPEB4 IP fraction, x-axis mean mRNA expression in Input and IGg control fractions, red and yellow points mark genes called significant by DESeq2 (for CPEB4 binding), n = 2 per fraction. **c,** Upper sub-panel, summed UMI counts for Cpeb genes w.r.t. cell types along the NSC lineage, lower sub-panel, quantification Cpeb genes in neurospheres derived from NSCs (aNSCs) by qPCR. **d**, De-novo motif analysis utilizing the *homer2* tool (Methods) in 3'UTRs that change along the NSC lineage, left result for *in vivo* and right for *in vivo* NSCs. **e,** Relative location of the CPE motif (from Fig. 3f) in 3'UTRs that get shorter or longer along the NSC lineage (x-axis, from Fig. 1d), y-axis, per 3'UTR: 0 = CPE is most proximal and 1 most distal, average if multiple CPEs per UTR, Wilcoxson rank sum test, two-sided. **f,** Comparison of proteomics (from Fig. 3) to CPEB4 RIP, density plot, x-axis fold-change in protein expression (aNSC vs. qNSC), groups colored in red if enriched in CPEB4 binding (CPEB4 substrate, from **b**) and grey depleted for CPEB4 binding (control gene set for low-CPEB4 binding), Wilcoxson rank sum test, two-sided. **g**, Differential expression of candidate genes in ASD vs. healthy controls, fold-changes from DESeq2, y-axis, per cell type, x-axis, log2-fold-change in gene expression, color code, p-value thresholds (FDR corrected) per cell-type. **h**, Expression trend of selected genes, upstream- and co-regulators of APA or ploy(A)-tail length (GO categories: GO:0006378 and GO:0031124), scRNA-seq from Fig. 1, total expression summed over single cells (x-axis) against correlation of expression vs. pseudotime (y-axis). **i.** Full membrane western blot used in this study (Figure 3j, right panel). **A:** Western blots showing the distribution of CPEB4 between the cytoplasmic (C) and nuclear (N) fractions in WT and Aplp1-/- mice with and without BMP4 treatment. T= Total protein. GAPDH and Histone H3 were used as cytoplasmic and nuclear markers.

*Extended Data Figure 5*

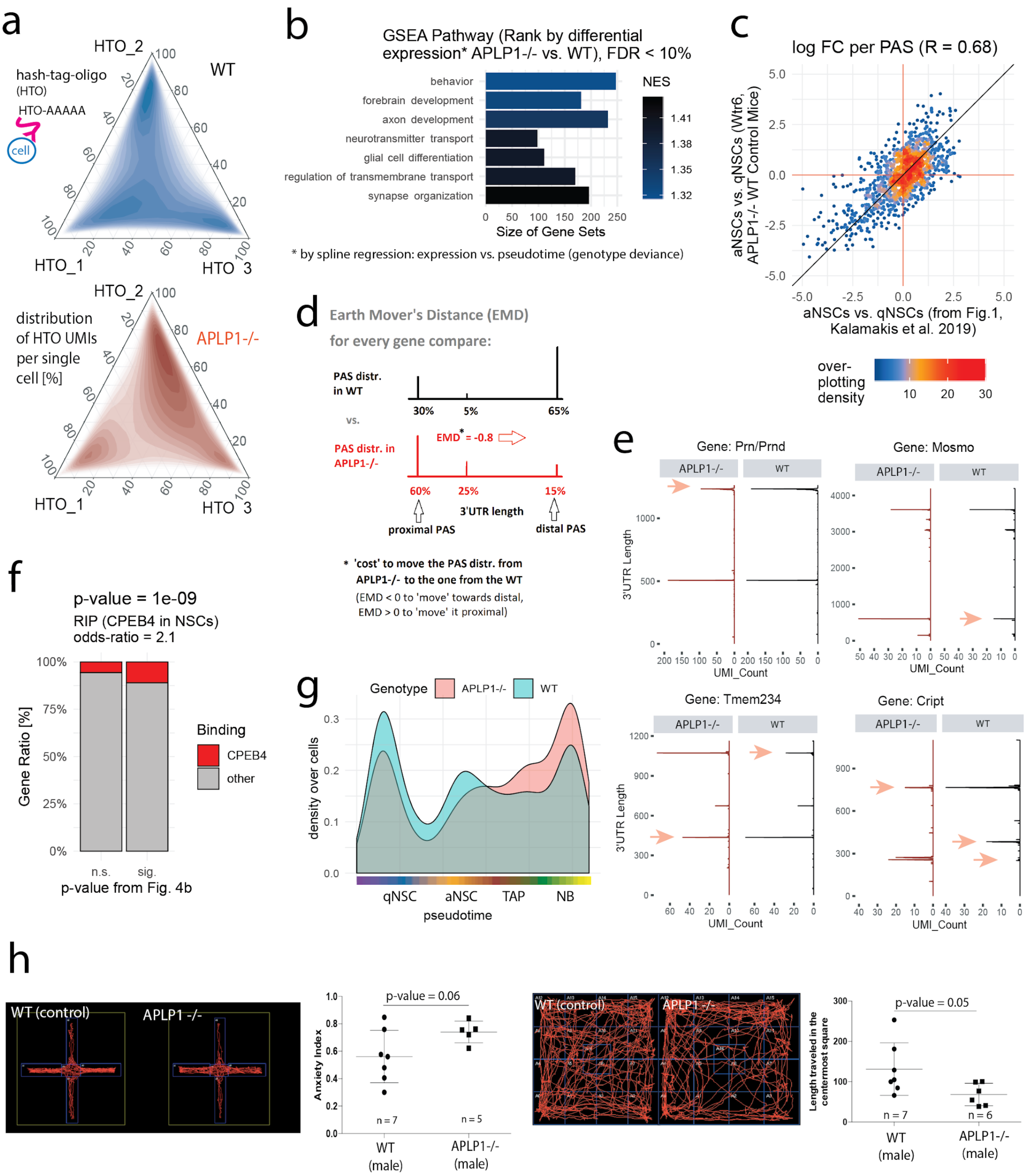

***Extended Data Figure 5* - Analysis of alterations in 3'UTRs comparing APLP1-/- to wildtype mice.**

**a,** Ternary plots for hash-tag-oligo (HTO) sequencing, upper sub-panel WT sample, lower sub-panel APLP1-/-, each HTO marks one biological replicate, relative distribution of all three HTOs per single cell, shown as local density (dark = high, bright = low). **b**, Gene set enrichment analysis, spline regression along pseudotime (from **g**) for differences in gene expression (APLP1-/- vs. WT), deviance explained by the genotype (ANODEV) used for gene ranking, high NES indicates association of gene category with genotype differences. **c,** Double-fold-change scatterplot, every point one PAS, direction: aNSCs to qNSCs, x-axis, scRNAseq shown in Fig. 1, y-axis, WT control mice for APLP1-/- scRNAseq run, agreement (high correlation) indicates reproducibility between sequencing batches in 3'UTR changes. **d**, Schematic depiction of the Earth Mover's distance (EMD) to quantify differential 3'UTR usage in APLP1-/- vs. wildtype (Fig. 4a), EMD is illustrated using one fictitious example gene. **e**, Differential 3'UTR distribution between APLP1-/- and wildtype, four example genes, 3' raw mapping position, UMIs summed over all cells per genotype (pseudo-bulk), arrows indicate differences, gets longer in APLP1-/-: Tmem234, gets shorter: Mosmo, Prn/Prnd and Cript. **f**, Intersection of CPEB4 RIP (Fig. 3h) and genes with 3'UTR alterations between APLP1-/- and wildtype (as in Fig. 4d), Fisher's exact test. **g**, scRNAseq for APLP-/- and wildtype control, pseudotime computed using *monocle2*, density over pseudotime shown per genotype. **h,** Further behavioral tests in APLP1-/- mice, upper panel, elevated plus maze (EPM) test to examine anxiety between APLP1-/- mice and control mice, left, example runs, red tracing lines of mouse location in the four arms of the EPM, lower panel, open filed test to examine anxiety in APLP1-/- mice vs. control mice, red tracing lines of mouse location in the open field (representative images), t-tests, two-sided, Welch's correction; error bars indicate standard deviation.

### Supplementary Tables

| Mapping 10X genomics (scRNAseq), paired-end, Bowtie2 for 3'peak calling | | | | |
|---|---|---|---|---|
| **Run** | **Sample_ID** | **Experimenter** | **Genotype** | **NSC isolation** |
| 1 | old_NSC | Georgios Kalamakis | WT | freshly, FACS |
| 2 | young_NSC | Georgios Kalamakis | WT | freshly, FACS |
| 3 | EGF_NSC | Nikhil George | WT | cultured |
| 4 | BMP4_NSC | Nikhil George | WT | cultured |
| 5 | APLP1_KO | Nikhil George | APLP1-/- | freshly, FACS |
| 6 | APLP1_WT | Nikhil George | WT | freshly, FACS |
| **Run** | **Mapping Rate** | **Number of mice** | **Number of cells** | **Hash-Tag-Oligos used** |
| 1 | 95.55% | 8 | 1716 | FALSE |
| 2 | 95.45% | 4 | 2035 | FALSE |
| 3 | 95.34% | 3 | 1410 | FALSE |
| 4 | 95.65% | 3 | 887 | FALSE |
| 5 | 96.04% | 3 | 1879 | TRUE |
| 6 | 96.17% | 3 | 2209 | TRUE |

**Table 1,** Information and summary about scRNAseq runs

| Symbol | Gene | log2FC | p.value | p.adjust | UMI_Count |
|---|---|---|---|---|---|
| Pmpcb | ENSMUSG00000029017 | -2,2303714 | 0,0002691 | 0,05133485 | 122 |
| Lypd6 | ENSMUSG00000050447 | -2,1250404 | 0,00025946 | 0,05133485 | 112 |
| Ostc | ENSMUSG00000041084 | -1,9466529 | 8,39E-05 | 0,02988535 | 142 |
| Tmem237 | ENSMUSG00000038079 | -1,9415733 | 0,00088712 | 0,09443402 | 129 |
| Sdc2 | ENSMUSG00000022261 | -1,9336141 | 0,00015456 | 0,04176934 | 175 |
| Cstf2 | ENSMUSG00000031256 | -1,8924245 | 0,00067408 | 0,08259773 | 103 |
| Phactr3 | ENSMUSG00000027525 | -1,8851021 | 0,0010254 | 0,09883928 | 122 |
| Rplp1 | ENSMUSG00000007892 | -1,8830506 | 9,22E-05 | 0,02988535 | 145 |
| Mpc1 | ENSMUSG00000023861 | -1,870334 | 0,00103624 | 0,09883928 | 121 |
| Selenos | ENSMUSG00000075701 | -1,8320586 | 1,74E-05 | 0,00806403 | 251 |
| Acadm | ENSMUSG00000062908 | -1,8237949 | 0,00093182 | 0,09443402 | 130 |
| 1110065P20Rik | ENSMUSG00000078570 | -1,7950665 | 0,00039176 | 0,07058175 | 165 |
| Pyroxd1 | ENSMUSG00000041671 | -1,7543004 | 0,00073702 | 0,08453656 | 137 |
| Ramp1 | ENSMUSG00000034353 | -1,6902439 | 0,00014196 | 0,04176934 | 282 |
| Slc39a12 | ENSMUSG00000036949 | -1,5383665 | 0,00021238 | 0,04919641 | 214 |
| Atp8a1 | ENSMUSG00000037685 | -1,5247445 | 0,00068248 | 0,08259773 | 214 |
| Ogdh | ENSMUSG00000020456 | -1,3901235 | 0,00068768 | 0,08259773 | 294 |
| Grm3 | ENSMUSG00000003974 | -1,3442839 | 1,00E-06 | 0,00108253 | 777 |
| Tril | ENSMUSG00000043496 | -1,3074792 | 3,60E-07 | 0,00058302 | 666 |
| Dbt | ENSMUSG00000000340 | -1,2736417 | 0,00075595 | 0,08453656 | 266 |
| Dynlrb1 | ENSMUSG00000047459 | -0,9868246 | 0,00025133 | 0,05133485 | 406 |
| Eno1b | ENSMUSG00000059040 | -0,8080414 | 0,00052833 | 0,07365145 | 1112 |
| Gnao1 | ENSMUSG00000031748 | -0,7410113 | 0,00049853 | 0,07365145 | 1537 |
| Usp22 | ENSMUSG00000042506 | 0,75960107 | 0,00054506 | 0,07365145 | 852 |
| Psip1 | ENSMUSG00000028484 | 1,27667282 | 1,71E-05 | 0,00806403 | 321 |
| Zfp451 | ENSMUSG00000042197 | 1,34205401 | 0,00091472 | 0,09443402 | 287 |
| Rlim | ENSMUSG00000056537 | 1,42289025 | 0,00042927 | 0,07326897 | 174 |
| Pnn | ENSMUSG00000020994 | 1,66244572 | 0,00050821 | 0,07365145 | 159 |
| Tpx2 | ENSMUSG00000027469 | 1,70641943 | 1,13E-05 | 0,00806403 | 209 |
| Ier2 | ENSMUSG00000053560 | 1,86860652 | 0,00051876 | 0,07365145 | 127 |
| Ccnb2 | ENSMUSG00000032218 | 1,89299362 | 1,62E-05 | 0,00806403 | 159 |
| Scrib | ENSMUSG00000022568 | 1,94105949 | 0,00021105 | 0,04919641 | 161 |
| Ubtf | ENSMUSG00000020923 | 2,045709 | 3,13E-05 | 0,01270397 | 125 |

**Table 2,** Differential expression in between APLP1-/- and wildtype, DESeq2 (pseudo-bulk approach = UMIs summed over biological replicates for each gene), 10% FDR.

Supplementary Images

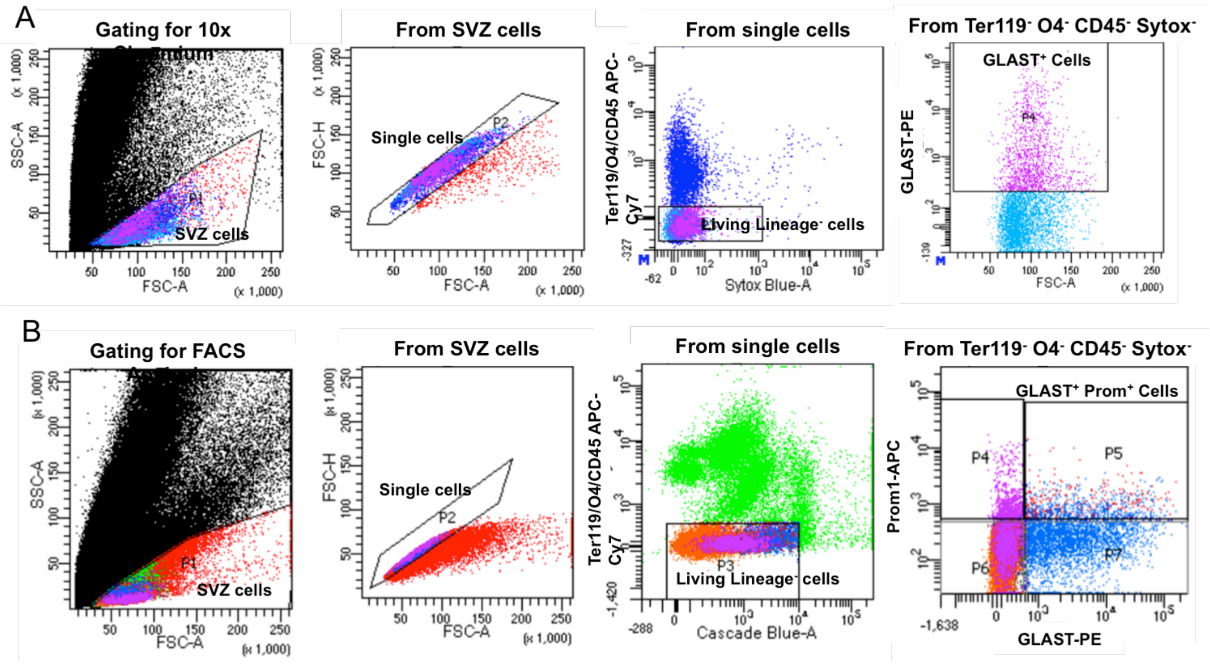

**FACS strategies used for vSVZ NSC lineage isolation and analysis:** A) Sorting strategy for the enrichment of GLAST+ cells used for 10x chromium scRNAseq approach. B) Gating strategy used for the FACS analysis of the number of NSCs from APLP1 -/- and WT mice.
